# Integrates JA and ethylene signals in defense against brown spot disease: the role of NaWRKY70

**DOI:** 10.1101/2023.02.01.526687

**Authors:** Na Song, Jinsong Wu

## Abstract

Production of phytoalexins scopoletin and scopolin is regulated by jasmonate (JA) and ethylene signals in *Nicotiana* species in response to *Alternaria alternata*, the necrotrophic fungal pathogen causing brown spot disease. However, how these two signals are coordinated to control this process remains unclear. Here, we found that levels of these two phytoalexins and transcripts of their key enzyme gene *feruloyl-CoA 6’-hydroxylase 1* (*NaF6’H1*) were synergistically induced in wild tobacco *N. attenuata* by co-treatments of methyl jasmonate (MeJA) and ethephon, but were not altered by treatments with either MeJA or ethephon along. Through co-expression and functional analysis we identified NaWRKY70 as the key regulator for this synergistic induction. NaWRKY70 not only directly bound to *NaF6’H1* promoter and activated its expression but also served as a regulation node to integrate JA and ethylene signals. Acting like locks, NaJAZe1, NaJAZe2 and NaEBF2 interacted with NaWRKY70 separately to suppress *NaF6’H1* expression. Meanwhile, NaMYC2b and NaEIN3-like 1functioned as transcription regulators of *NaWRKY70* and activated of *NaF6’H1* expression by protein interaction with NaWRKY70. Finally, NaWRKY70 controlled JA-Ile production by binding and activating its biosynthetic genes. Thus, our data uncovers a novel but complicate regulation network of phytoalexins by two phytohormonal signals, and demonstrates that NaWRKY70 integrates both JA and ethylene signals through transcriptional regulation and protein interaction to regulate scopoletin and scopolin biosynthesis and plant resistance to *A. alternata*.

## Introduction

To adapt to the complicated and dynamic environments, plants produce hundreds of thousands of specialized secondary metabolites. Many of them are responsible for growth (Erb and Kliebenstein 2020), development (Vaughan *et al*., 2015) and plant immunity (Ahuja *et al*., 2012; Piasecka *et al*., 2015). Phytoalexins, a group of low molecular mass secondary metabolites elicited after pathogen attack or elicitors, are important “chemical weapons” for plant resistance to pathogen (Pedras *et al*., 2011; Ahuja *et al*., 2012; Wu 2020). In *Arabidopsis*, camalexin is a major phytoalexin for resistance to necrotrophic pathogens such as *Botrytis cine*rea (Ferrari *et al*., 2007), *Alternaria brassicicola* (Nafisi *et al*., 2007), *Sclerotinia sclerotiorum* (Stotz *et al*., 2011) and *Phytophthora brassicae* (Schlaeppi *et al*., 2010). Capsidiol and scopoletin are two major phytoalexins produced in *Nicotiana* species in response to pathogens (El Oirdi *et al*., 2010; Grosskinsky *et al*., 2011; Sun *et al*., 2014b; Song *et al*., 2019).

*Alternaria alternata* (tobacco pathotype), a necrotrophic fungal pathogen, is the causative agent of brown spot disease in *Nicotiana* species including *Nicotiana tabacum* (LaMondia 2001) and wild tobacco *Nicotiana attenuata* (Schuck *et al*., 2014; Sun *et al*., 2014a). Previously, we have demonstrated that capsidiol, scopoletin and scopolin are three conserved major phytoalexins produced in *Nicotiana* species in response to *A. alternata* (Sun *et al*., 2014b; Li and Wu 2016; Song *et al*., 2019; Long *et al*., 2021). Scopoletin, a 7-hydroxy-6- methoxy-phenolic coumarin, is biosynthesized through phenylpropanoid pathway (Kai *et al*., 2006). Mutant analysis revealed that feruloyl-CoA 6’-hydroxylase 1 (NaF6’H1) was the key enzyme for the formation of scopoletin (Kai *et al*., 2008). Scopolin, a β-glycoside form of scopoletin, also acts as a phytoalexin against *Tobacco mosaic virus* and *A. alternata* (Chong *et al*., 2002; Li and Wu 2016).

Jasmonate (JA) and ethylene are two major plant hormones involved in defense against necrotrophic fungi and herbivores (McDowell and Dangl 2000; Onkokesung *et al*., 2010). In response to pathogen or herbivore attack, plants synthesize JA-isoleucine (JA-Ile), the active form of jasmonate, and ethylene. JA-Ile can combine with complex receptor SCF^COI1^, which ubiquitinate repressors JASMONATE ZIM DOMAIN (JAZ) proteins to release regulators, including MYC2, MYC3 and MYC4, thereby activating downstream response genes (Xu *et al*., 2002; Thines *et al*., 2007; Katsir *et al*., 2008; Sheard *et al*., 2010). JAZs also interact with several other transcription factors to regulate JA-mediated responses (Lv *et al*., 2017; Wang *et al*., 2020; Chen *et al*., 2021). Ethylene is perceived by a series of receptors, which combine with the CTR1, a Raf-like kinase constitutive ethylene response1 (Guo and Ecker 2004). The essential positive regulator EIN2 (ETHYLENE INSENSITIVE 2) acts the downstream of CTR1. In the presence of ethylene, CTR1 is inactive, resulting in the accumulation of transcription factors EIN3 and EIN3-like1 (EIL1), subsequently triggers various ethylene response genes (Yanagisawa *et al*., 2003; Ju *et al*., 2012). The stabilization of EIN3 and EIL1 are dependent on two EIN3 binding F-box proteins EBF1 and EBF2, which are the master repressors of ethylene signaling pathway (Yang *et al*., 2015; Zhao *et al*., 2021). Previously, we have demonstrated that scopoletin and scopolin are two important phytoalexins required for *A. alternata* resistance, and their production is strongly impaired in transgenic plants silenced with *NaAOC* or *NaACO*, two key enzyme genes for JA and ethylene biosynthesis (Sun *et al*., 2014b; Li and Wu 2016; Sun *et al*., 2017), suggesting both JA and ethylene signals are crucial for biosynthesis of scopoletin and scopolin. However, the detail mechanism how these two phytoalexins are coordinately regulated by JA and ethylene signals remains unclear.

WRKY transcription factors, a large family of regulatory proteins only present in plants, are involved in growth, development, defense against pathogen, and response to external stimuli (Eulgem *et al*., 2000; Rushton *et al*., 2010; Chen *et al*., 2012; Schluttenhofer and Yuan 2015). They are characterized by the conserved WRKYGQK and zinc finger-like motifs, recognizing cis-elements W-boxes in the promoter of target genes and activating or inhibiting their expression (Rushton *et al*., 2010; Chen *et al*., 2012). In *Arabidopsis*, *AtWRKY33* acts as a master transcription factor to directly regulate biosynthesis genes *PAD3* and *CYP71A13* expression for camalexin to defense against *B. cinerea* (Mao *et al*., 2011). Recently, it was found that CPK5/CPK6 and MPK3/MPK6-mediated the differential phosphorylation of AtWRKY33 cooperatively participated in camalexin biosynthesis (Zhou *et al*., 2020). Interestingly, analysis of *NaF6’H1* promoter revealed that its expression was likely regulated by some unknown WRKYs as five W-boxes occurred in the promoter region. Thus, it would be very interesting to investigate whether any WRKYs can integrate JA and ethylene signals to regulate scopoletin and scopolin biosynthesis.

Here, we showed that levels of phytoalexins, scopoletin and scopolin, and expression of their key enzyme gene *NaF6’H1* were remarkably and synergistically induced in *N. attenuata* by JA and ethylene signals. Through co-expression and functional analysis, we identified NaWRKY70 as a key regulator responsible for this synergistic induction. NaWRKY70 could directly bind to *NaF6’H1* promoter and activate its expression. Furthermore, we uncovered the mechanism of this synergistic induction of scopoletin and scopolin by JA and ethylene. Repressors of JA and ethylene signaling, NaJAZe1, NaJAZe2 and NaEBF2, could interact with NaWRKY70 to inhibit *NaF6’H1* expression. In addition, NaMYC2b and NaEIN3-like 1 were two key regulators of JA and ethylene pathways; both were required for *NaWRKY70* expression, and interacted with NaWRKY70 protein to activate *NaF6’H1* expression. Finally, NaWRKY70 also controlled JA-Ile biosynthesis by binding and activating JA biosynthetic genes as a feedback. Thus, our work extends our understanding of synergistic induction of phytoalexins, and provides new insight into defense responses of *Nicotiana* plants after *A. alternata* attack.

## Results

### Synergistic induction of phytoalexins scopoletin and scopolin by JA and ethylene signals

In response to *A. alternata*, *N. attenuata* plants activated both JA and ethylene signaling, and both of which are required for the biosynthesis of phytoalexins scopoletin and its β-glycoside form, scopolin (Figure 1A and 1B; Sun *et al*., 2014b; Li and Wu 2016; Sun *et al*., 2017). Interestingly, these two phytoalexins could not be induced in leaves treated with either methyl jasmonate (MeJA) or ethephon (an ethylene-releasing agent) alone. These results led us hypothesize that scopoletin and scopolin might be only induced by activation of both JA and ethylene signals. We thus investigated scopoletin and scopolin accumulation by exogenous co-treatments with MeJA (1 mM) and ethephon (5 mM). Indeed, strong blue fluorescence, a signature of scopoletin and scopolin under UV light, were observed on lamina of source-sink transition leaves (0 leaves) after co-treatments with MeJA and ethephon for 3 days, whereas no blue fluorescence were accumulated in leaves treated with MeJA or ethephon alone (Figure 1C).

**Figure 1.**
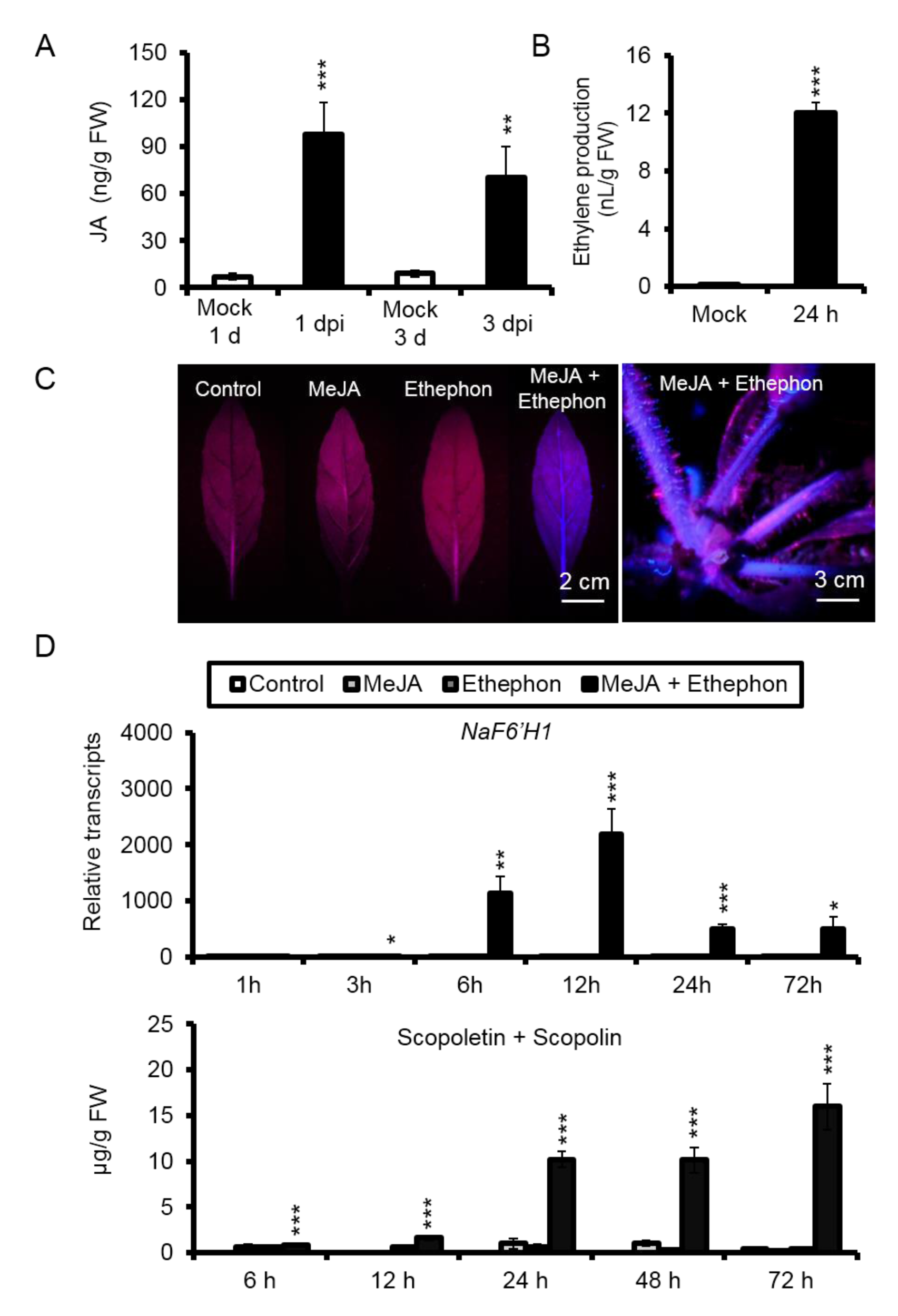
Synergistic induction of phytoalexins scopoletin and scopolin by JA and ethylene signals. **(A)** Mean (±SE) *A. alternata*-induced JA levels were measured by HPLC-MS/MS in six biological replicates of source-sink transition (0) leaves of WT plants at 1 and 3 dpi. **(B)** A. alternata-induced ethylene production for 24 h was measured by GC-MS in four biological replicates of 0 leaves at 1 dpi. **(C)** Photos of 0 leaves and whole WT plants under UV after 3 d treatments of water control, 50 mM MeJA, 1 mM ethephon, and co-treatments of MeJA and ethephon. The blue fluorescence of rosette 0 leaves (left) and whole plant (right) were obvious under the UV-light when treated with MeJA and ethephon simultaneously. **(D)** Mean (±SE) relative expression levels of NaF6’H1, and scopoletin and scopolin levels were measured in five biological replicates of 0 leaves at 1, 3, 6, 9, 12, 24 and 72 h after treatments of water control, 50 mM MeJA, 1 mM ethephon, and co-treatments of MeJA and ethephon. Asterisks indicate the levels of significant differences between control and treated samples with the same time points (Student’s t-test: *, P<0.05; **, P<0.01; ***, P<0.005).

We next measured the transcripts of *feruloyl-CoA 6’-hydroxylase 1* (*NaF6’H1*), the key enzyme gene for scopoletin biosynthesis by quantitative RT-PCR (qRT-PCR). When compared to water control treatments, *NaF6’H1* transcripts were strongly elicited up to 600-fold in 0 leaves treated with MeJA and Ethephon simultaneously for 6 h, peaked at 2189-fold for 12 h, and gradually decreased to around 500-fold from 24 h to 72 h (Figure 1D). However, *NaF6’H1* transcripts were not altered in 0 leaves when treated with MeJA or ethephon alone (Figure 1D). Consistently, when leaves were treated with MeJA and ethephon simultaneously, levels of scopoletin and scopolin increased significantly at 6 h, and peaked 15.97 ± 2.53 µg g^-1^ fresh leaves at 72 h, while phytoalexins level did not differ between leaves treated with water and MeJA or ethephon only (Figure 1D). In addition, co-treatments with MeJA and ethephon also elevated production of scopoletin and scopolin in the lamina of mature leaves, mid-rib of young and mature leaves (Supplementary Fig. S1).

Interestingly, the synergistic induction of scopoletin and scopolin by MeJA and ethylene seemed to be only conserved in the genus of *Nicotiana*. We also observed it in cultivated tobacco K326 (Wu and Song 2021), but not in *Arabidopsis thaliana*, *Gossypium hirsutum,* and Solanaceae plants *Solanum tuberosum* and *Solanum lycopersicum* (Supplementary Fig. S2).

To further explore the molecular mechanism underlying how JA and ethylene signals regulated the biosynthesis of scopoletin and scopolin, we detected the transcriptional levels of *NaF6’H1* and scopoletin and scopolin production in WT and transgenic plants silenced with *NaCOI1* (JA-insensitive, irCOIl), *NaAOC* (deficient in JA biosynthesis; irAOC), *NaACO* (ethylene reduced; irACO), or over-expressed with *Arabidopsis etr1* (ethylene insensitive; Ov-etr1). Our results indicated that the synergistic induction of both *NaF6’H1* transcripts and phytoalexins production by MeJA and ethephon were observed in WT and JA-deficient irAOC leaves, but were abolished in JA-insensitive irCOI1 leaves (Supplementary Fig. S3A and 3B). Similarly, no effect of MeJA and ethephon co-treatments on the expression of *NaF6’H1* and scopoletin accumulation were observed in ethylene-insensitive Ov-etr1 plants (Supplementary Fig. S3C and 3D).

Taken together, we observed the phenomena of synergistic induction of *NaF6’H1* expression and scopoletin and scopolin production by MeJA and ethephon, and this specific induction required both endogenous intact JA and ethylene signaling pathways.

### Screening the key transcription factors involved in the synergistic induction of scopoletin and scopolin by JA and ethylene signals

To identify the key transcription factors involved in the synergistic induction of phytoalexins by JA and ethylene signals, we performed an transcriptome analysis with RNA sequencing in 0 leaves of WT plants with 6 h treatments of water control, MeJA (1 mM), ethephon (5 mM), or both MeJA and ethephon. Co-expression analysis revealed that 132 transcription factors (TFs) showed synergistic induction of their expressions by MeJA and ethephon co-treatments similar to *NaF6’H1*, and 16 candidate TFs were enriched (log2 (fold change)≥3 when compared with MeJA o ethephon alone; Supplementary Table 2).

We next silenced these 16 TFs individually via virus-induced gene silencing to identify those involved in scopoletin and scopolin biosynthesis, including NaMYC4-like, NaMYC2-like, NaMYB24-like, NaMYB57-like, NaHAT5-like, NabHLH18-like, NaERF17-like, NaZinc655-like, NaWRKY70, NaMYB4-like, NaWRKY75-like, NaWRKY70-like, NaWRKY41-like, NabHLH137-like, NabHLH93-like and NaMYB44-like. Our results indicated that *A. alternata*-elicited *NaF6’H1* expression were significantly reduced in NaMYC4-, NaWRKY70-, NaWRKY75-like-, and NaMYB44-like-silenced plants (Supplementary Fig. S4).

Thus, NaWRKY70 was selected as a candidate for further analysis in this study.

### *NaWRKY70* expression is induced by *A. alternata* in JA/ethylene-, and age-dependent ways

NaWRKY70 encoded a peptide of 225 amino acids, containing a conserved WRKYGQK domain and a Cx7Cx23HxC zinc-finger (Supplementary Fig. S5A). When *NaWRKY70*::*eGFP*, which was driven by the CaMV 35s promoter, was transformed into protoplasts of *N. attenuata*, strong GFP fluorescence was observed in the nucleus (Supplementary Fig. S5B). Phylogenetic tree of WRKYs from *Arabidopsis*, *Solanaceae* and other plants, was constructed by the neighbor-joining (NJ) program. NaWRKY70, NtWRKY70 and NtWRKY4 were clustered together (Supplementary Fig. S5C). These results confirmed that NaWRKY70 was a member of WRKY family, and localized in the nucleus.

We performed real-time PCR to investigated *NaWRKY70* expression in WT, irAOC, irCOI1, irACO and Ov-etr1 plants. As expected, *NaWRKY70* transcripts were strongly elicited and increased gradually in leaves co-treatment of MeJA and ethephon at 1, 3 and 6 h, but did not differ in leaves treated with MeJA or ethephon alone at all three time points (Figure 2A). This synergistic induction of *NaWRKY70* by MeJA and ethephon was reproduced in JA-deficient irAOC and ethylene-reduced irACO leaves, but was abolished in JA-insensitive irCOI1 or ethylene-insensitive Ov-etr1 plants (Figure 2B).

**Figure 2.**
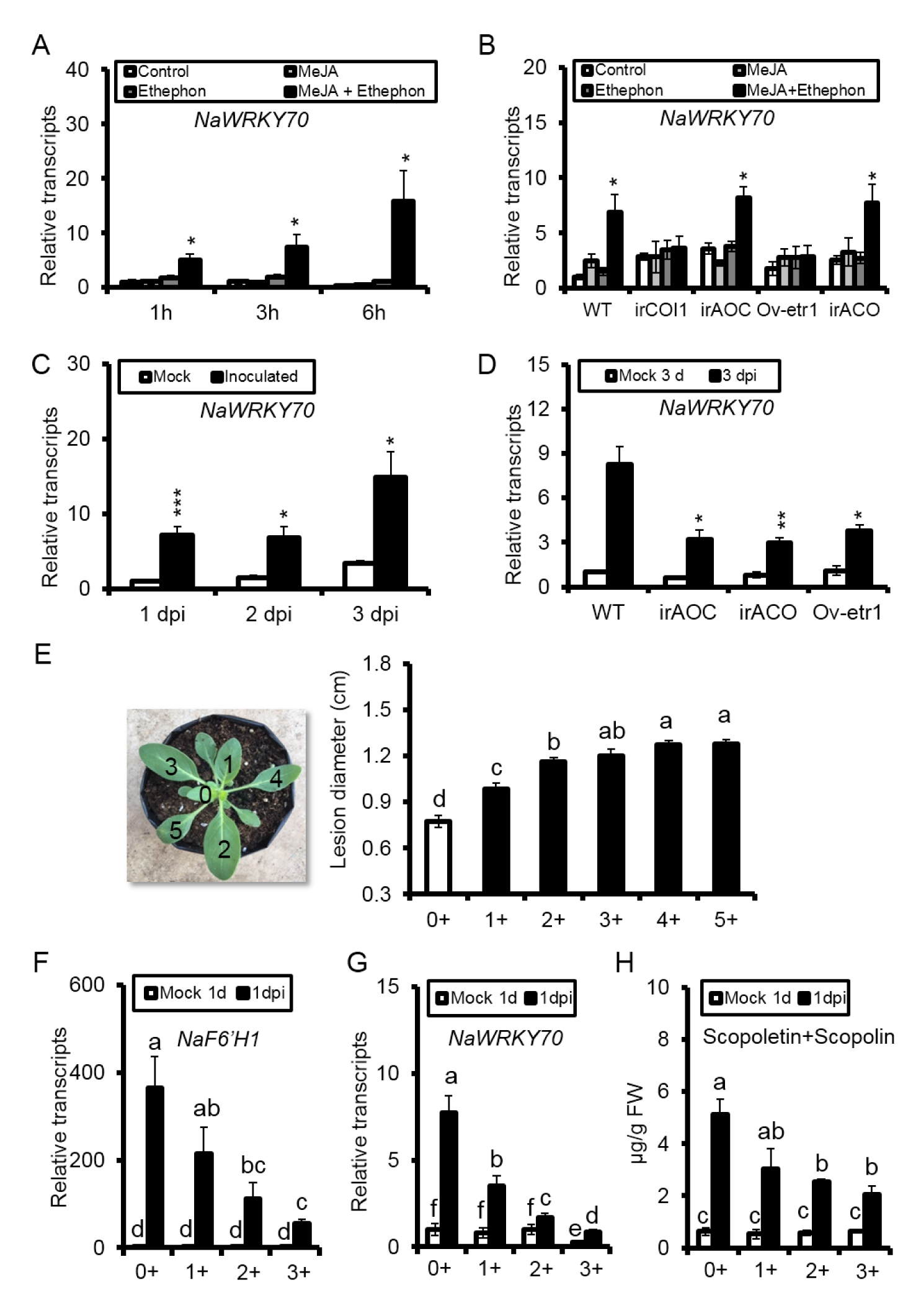
NaWRKY70 expression is induced by A. alternata in JA/ethylene-, and age-dependent ways. **(A)** Mean (±SE) relative *NaWRKY70* expression levels were quantified in five biological replicates of 0 leaves by RT-qPCR at 1, 3 and 6 h in response to water, MeJA, ethephon, and co-treatments of MeJA and ethephon. Asterisks indicate the levels of significant differences between control and co-treated (MeJA and ethephon) samples (Student’s t-test: *, P<0.05). **(B)** Mean (±SE) relative expression levels of NaWRKY70 in five biological replicates of 0 leaves of WT, irAOC, irACO and Ov-etr1 plants after water, MeJA, ethephon, and co-treatment of MeJA and ethephon at 6 h. Asterisks indicate the levels of significant differences between control and MeJA and ethephon co-treated samples (Student’s t-test: *, P<0.05). **(C)** Mean (±SE) relative NaWRKY70 expression levels were analyzed in five-biological-replicated 0 leaves of WT after infection by A. alternata at 1, 2 and 3 dpi. Asterisks indicate the level of significant differences between mock and inoculated samples (Student’s t-test: *, P<0.05; ***, P<0.005). **(D)** Mean (±SE) relative expression levels of NaWRKY70 were measured in five replicates of 0 leaves WT, irAOC, irACO and Ov-etr1 plants at 3 dpi. Asterisks indicate the level of significant differences between WT and irAOC, irACO or Ov-etr1 plants after infection by A. alternata at 3 dpi (Student’s t-test: *, P<0.05; **, P<0.01; ***, P<0.005). **(E)** Mean (± SE) diameter of necrotic lesions of 15 biological replicates in different numbered rosette leaves of WT infected with A. alternata for 5 d (right). Numbering of the leaves at different phyllotaxic positions (nodes) on a rosette-stage plant was following Sun et al., (2014a). The leaf at node 0 (0+) is at the stage of source-sink transition, one phyllotaxic position younger than the first fully expanded leaf (1+). Similarly, the leaf at node 3 (3+) is also one phyllotaxic position younger than the leaf at node 4 (4+). Mean (±SE) relative expression levels of NaF6’H1 **(F)** and NaWRKY70 **(G)**, and scopoletin and scopolin levels **(H)** were measured in 8 biological replicates of 0 leaves of WT plants at 1 dpi. Different letters indicate significant differences by two-way ANOVA followed by Duncan’s test (P<0.05).

The transcriptional levels of *NaWRKY70* were also significantly increased in 0 leaves of WT plants after *A. alternata* inoculation at 1, 2 and 3 days (dpi) respectively (Figure 2C), and this *A. alternata*-induced expression were decreased by 50%, 70% and 60% in irAOC, irACO and Ov-etr1 plants respectively (Figure 2D), indicating that both JA and ethylene signaling pathways are required for *A. alternata*-induced *NaWRKY70* expression.

*A. alternata*, a necrotrophic fungal pathogen, causes brown spot disease in *Nicotiana* species. Interestingly, this disease usually occurs in mature leaves in both *N. tabacum* and *N. attenuata*, but hardly detect in young leaves (Cheng and Sun 2001; Sun *et al*., 2014b). Indeed, we also found that *N. attenuata* leaves became more susceptible to *A. alternata* with the increasing maturity of leaves in this study (Figure 2E). This phenomena was associated with higher *NaF6’H1* expression and scopoletin and scopolin production in young leaves (Figure 2F and 2H). Importantly, *NaWRKY70* was also highly expressed in young leaves, but decreasing while leaves become mature as *NaF6’H1* expression and scopoletin and scopolin production (Figure 2G). These results suggested that NaWRKY70 might account for the age-dependent susceptible to *A. alternata*.

### NaWRKY70 is a key transcription factor involved in the synergistic induction of scopoletin and scopolin by JA and ethylene signals, and *A. alternata* resistance

To further elucidate the role of NaWRKY70 in *A. alternata* resistance, stable transgenic *NaWRKY70*-silenced lines were generated by RNA interference (RNAi) through *Agrobacteria*-mediated transformation. We selected two independent, homozygous T2 lines of NaWRKY70-RNAi (1^#^ and 4^#^) for further analysis. *A. alternata*-elicited *NaWRKY70* expression in 0 leaves at 3 dpi was successfully silenced in both RNAi lines, with decreased by 95% and 98% respectively (Figure 3A). Importantly, *A. alternata*-elicited *NaF6’H1* transcripts were also reduced by more than 80%, which resulted 80% to 90% reduction of scopoletin and scopolin levels at 3 dpi (Figure 3B and 3C). We also found that *NaCCoAOMTs,* key enzyme genes of scopoletin biosynthesis, were showed to be down-regulated in both two NaWRKY70-RNAi lines (Supplementary Fig. S6). Moreover, NaWRKY70-RNAi plants were more susceptible to *A. alternata* than WT, as much bigger lesions were developed (Figure 3D). Finally, synergistic induction of *NaF6’H1*, scopoletin and scopolin by co-treatments of MeJA and ethephon was strongly impaired in 0 leaves of NaWRKY70-RNAi plants (Figure 3E-G), indicating that NaWRKY70 was the key transcription factor involved in the synergistic induction of scopoletin and scopolin by JA and ethylene signals, and *A. alternata* resistance.

**Figure 3.**
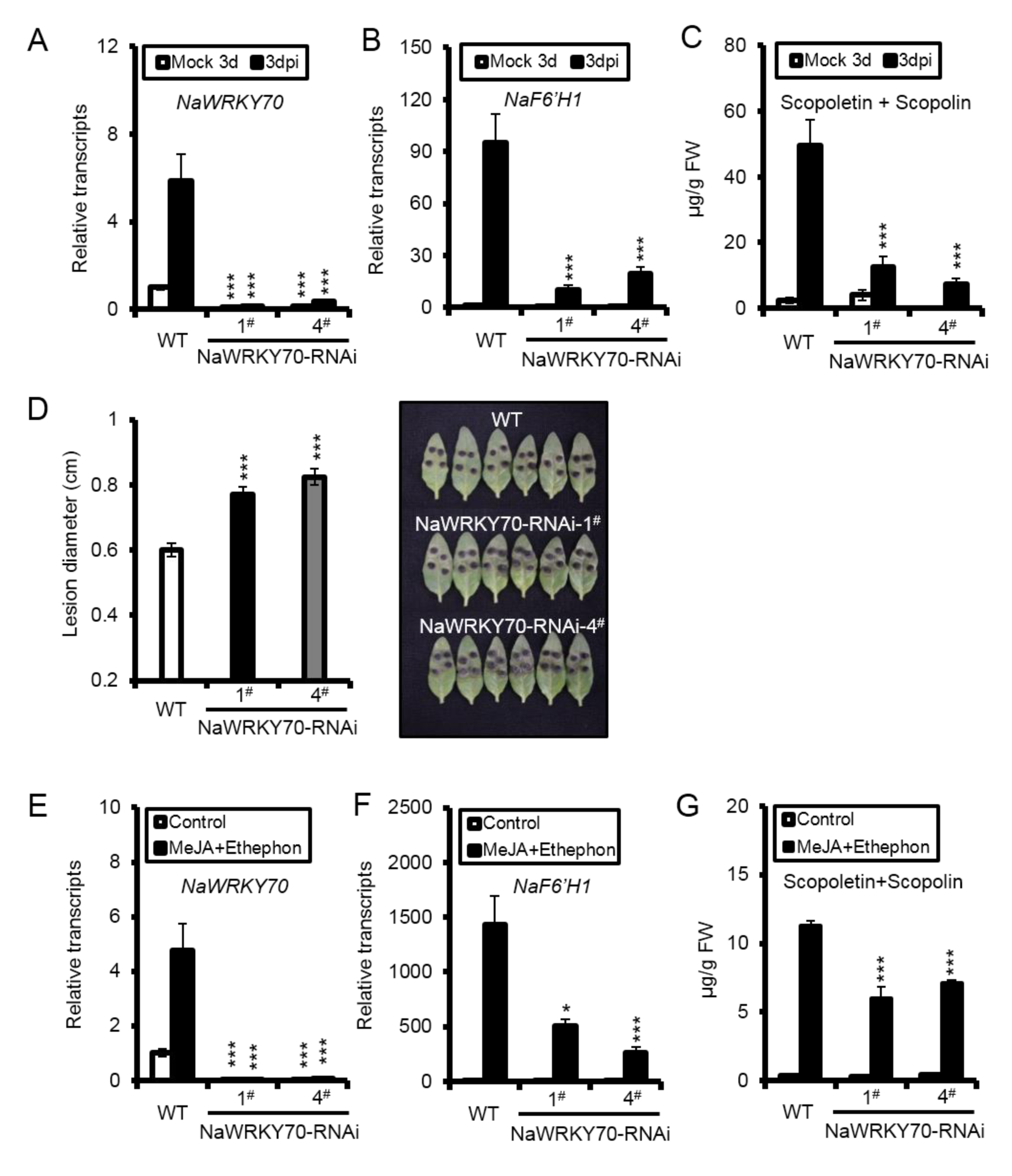
Silencing NaWRKY70 strongly impairs A. alternata (or co-treated with MeJA and ethephon)-induced NaF6’H1 expression, scopoletin and scopolin production. Mean (±SE) relative A. alternata-induced transcriptional levels of NaWRKY70 **(A)** and NaF6’H1 **(B)**, and scopoletin and scopolin levels **(C)** were measured in five biological replicates of 0 leaves of WT and two independent NaWRKY70 silencing transgenic plants (NaWRKY70-RNAi-1# and 4#) at 3 dpi. **(D)** Mean (± SE) diameter of necrotic lesions of 15 biological replicates of 0 leaves of WT, NaWRKY70-RNAi-1# and 4# infected with A. alternata for 5 d (left). Photos of 6 representative leaves of each different genotype at 5 dpi were taken (right). Mean (±SE) relative transcripts of NaWRKY70 **(E)** and NaF6’H1 **(F)**, and scopoletin and scopolin levels **(G)** were measured in five biological replicates of 0 leaves of different genotypes after MeJA and ethephon co-treatments. Asterisks indicate the levels of significant differences between WT and two NaWRKY70-RNAi lines with the same treatments (Student’s t-test: *, P<0.05; ***, P<0.005).

To further confirm the results of RNAi lines, we also generated two *NaWRKY70* knock-out mutants (*NaWRKY70*-cas-12^#^ and *NaWRKY70*-cas-13^#^) by CRISPR/Cas9 system. These two mutants contained different insertions at target sites.

*NaWRKY70*-cas-12^#^ had a T insertion in target 1 and an A insertion in target 2, while *NaWRKY70*-cas-13^#^ had a T insertion in target 1 and another T insertion in target 2 (Figure 4A). Similar to results of RNAi lines, both *A. alternata*-induced *NaF6’H1* and production of scopoletin and scopolin were also strongly impaired in *NaWRKY70* mutants (Figure 4B and 4C).

**Figure 4.**
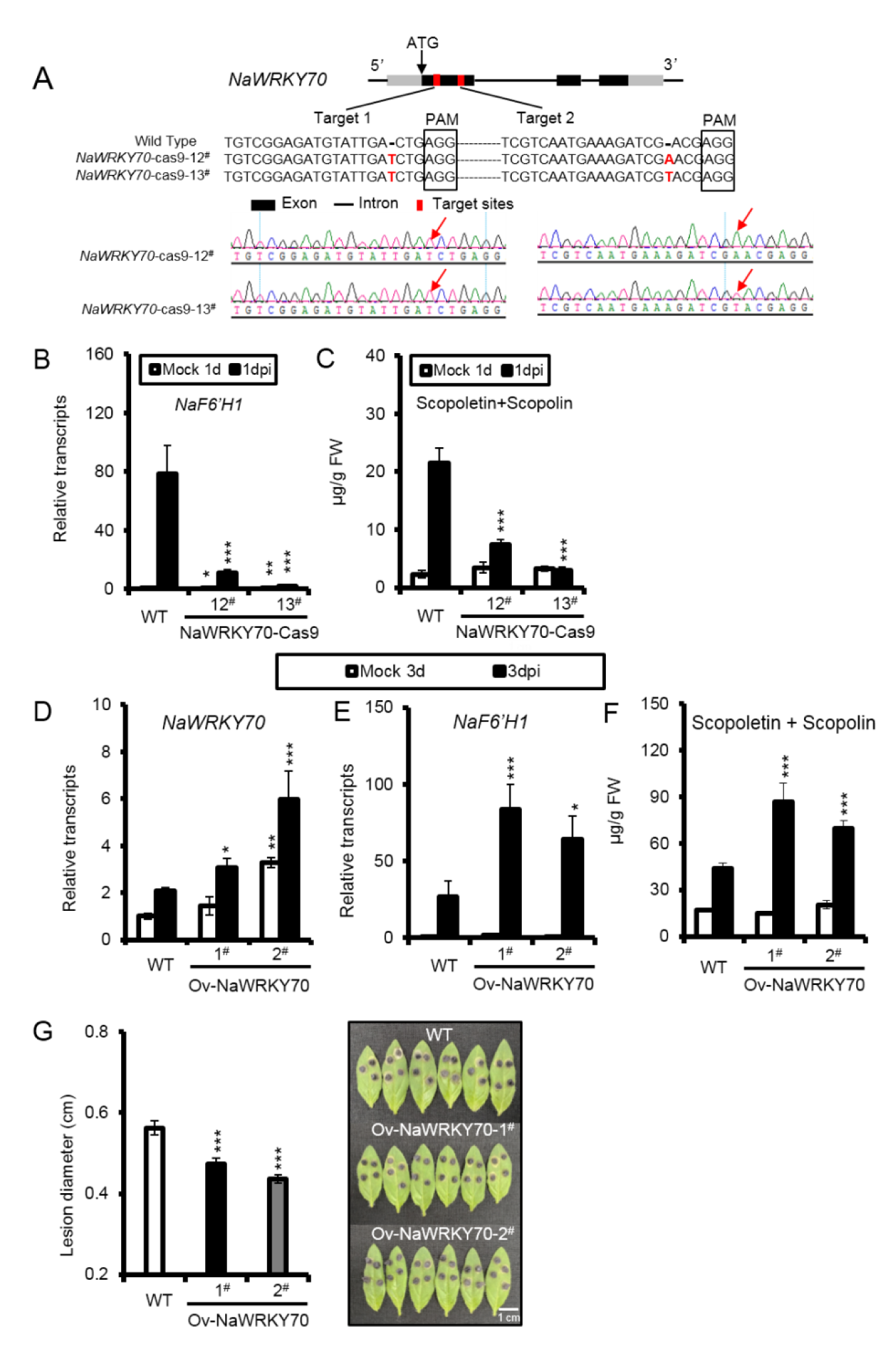
A. alternata-induced NaF6’H1 expression and scopoletin and scopolin biosynthesis in NaWRKY70 knock-out mutants and over-expression lines. **(A)** Schematic diagrams of NaWRKY 70 gene showing the mutation sites created by CRISPR/Cas9. Two sgRNA sequences of specific targets of NaWRKY70 were showed in details, thus generating NaWRKY70-cas9-12# and NaWRKY70-cas9-13# mutants. PAM sites were indicated by black squares. The insertion sites of target 1 and target 2 were indicated by red arrows. Mean (±SE) relative A. alternata-elicited NaF6’H1 expression levels **(B)** and A. alternata-induced scopoletin and scopolin levels **(C)** were measured in five biological replicates of 0 leaves of WT and two NaWRKY70 knock-out mutants (NaWRKY70-cas-12# and NaWRKY70-cas-13#) at 1 dpi. Asterisks indicate the levels of significant differences between WT and two NaWRKY70 mutants (Student’s t-test: *, P<0.05; **, P<0.01; ***, P<0.005). Mean (±SE) relative transcripts of NaWRKY70 **(D)** and NaF6’H1 **(E)**, and scopoletin and scopolin levels **(F)** were measured in five replicates of 0 leaves of WT and two stable transgenic NaWRKY70 over-expression (Ov-NaWRKY70-1# and 2#) plants at 3 dpi. **(G)** Mean (± SE) diameter of necrotic lesions of 15 biological replicates of 0 leaves of WT, Ov-NaWRKY70-1# and 2# after inoculation with A. alternata for 5 d (left). Photos of 6 representative leaves of each different genotype at 5 dpi were taken (right). Asterisks indicate the levels of significant differences between WT and two Ov-NaWRKY70-1# and 2# plants (Student’s t-test: *, P<0.05; **, P<0.01; ***, P<0.005).

We further generated stable transgenic *NaWRKY70* over-expression (Ov-NaWRKY70-1^#^and2^#^) plants through *Agrobacteria*-mediated transformation. Both the basal and induced levels of *NaWRKY70* were substantially increased, with 149% and 289% of that of WT at 3 dpi (Figure 4D). Concomitantly, *A. alternata*-induced expression levels of *NaF6’H1* in the Ov-NaWRKY70 lines were increased to 3.2- and 2.42-fold of that of WT respectively, and levels of both phytoalexins and *A. alternata* resistance were dramatically increased in Ov-NaWRKY70 lines compared with that of WT (Figure 4E-G).

All these results support the great role of NaWRKY70 in the synergistic regulation of phytoalexins scopoletin and scopolin by JA and ethylene signaling pathways, and *A. alternata* resistance.

### NaWRKY70 regulates scopoletin biosynthesis by directly binding and activating *NaF6’H1* promoter

As *NaWRKY70* was required for *A. alternata*-induced *NaF6’H1* expression and scopoletin accumulation, we hypothesized that the WRKY70 protein might bind directly to *NaF6’H1* promoter. *NaF6’H1* had five conserved W-boxes in its promoter region which could be potential binding sites of WRKYs. We thus designed biotin labeled probes containing these five W-boxes respectively for electrophoretic mobility shift assays (EMSA). NaWRKY70-GST protein were expressed and purified from *Escherichia coli*. EMSA indicated that NaWRKY70 protein could only bind to one of these five probes, probe 1 (Supplementary Fig. S7 and Figure 5A). This binding was specific, as the binding was attenuated by adding 50 times unlabeled cold probe, and abolished by 200 times cold probe. Importantly, the mutated probe lost the binding activity with NaWRKY70 (Figure 5A).

**Figure 5.**
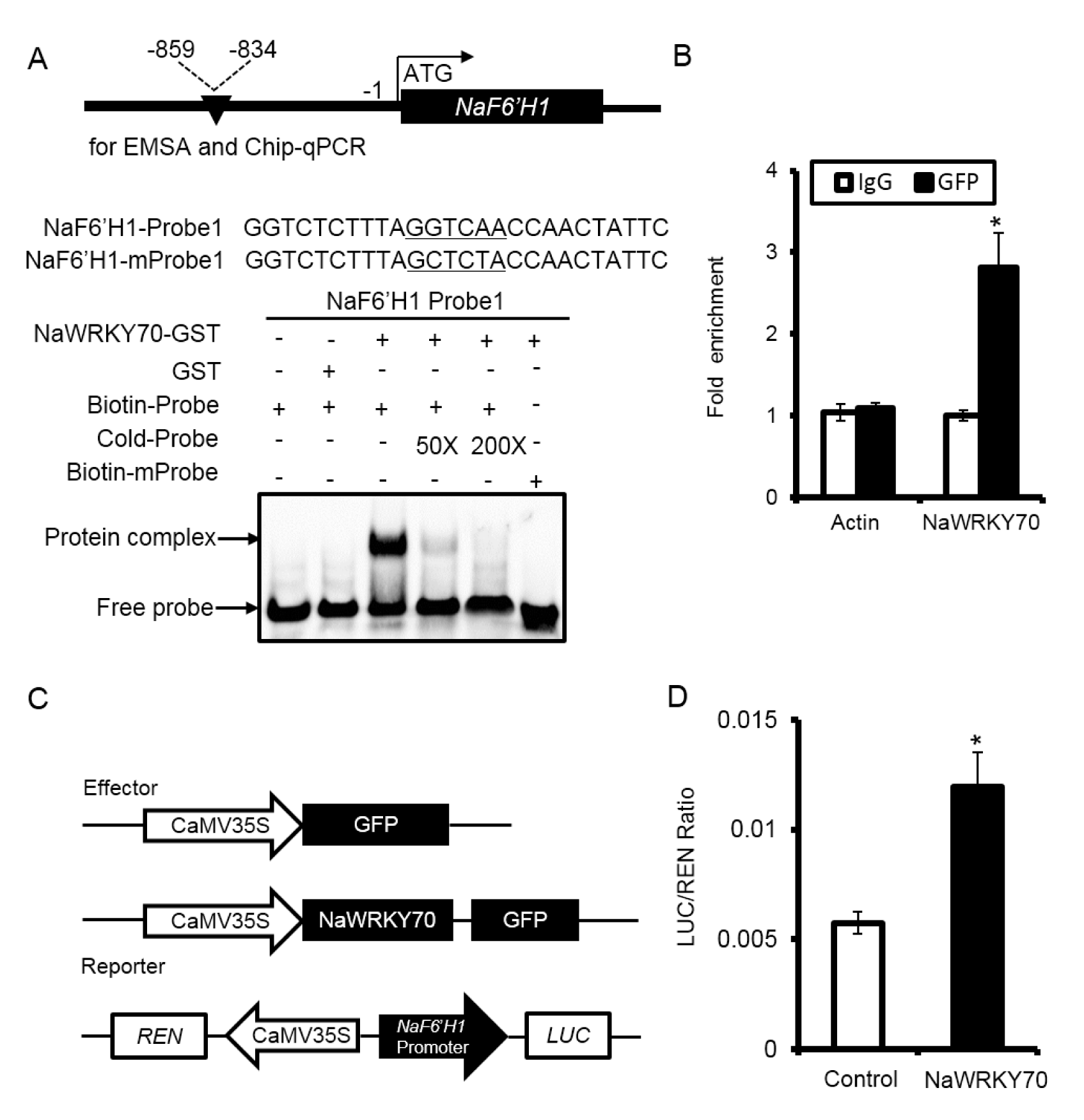
NaWRKY70 directly binds to the promoter of NaF6’H1 and activates its transcriptional activity. **(A)** Schematic diagrams of NaF6’H1 promoter and EMSA result of the binding of NaWRKY70 to NaF6’H1 promoter. The sequence and position of probe 1 (and mutated one: mProbe 1) in NaF6’H1 promoter were indicated. EMSA demonstrated that NaWRKY70 protein could specifically bind to probe 1. The mobility shift was abolished by adding cold unlabeled probes, and no signals were detected when mutant probe was used. **(B)** Detection of in vivo binding of NaWRKY70 to NaF6’H1 promoter by Chip-qPCR. In NaWRKY70-eGFP transgenic lines, higher levels of NaF6’H1 promoter regions around probe 1 were enriched with GFP antibody. Negative controls were imuno-precipitated with IgG or with GFP antibody but using primers detecting Actin gene. Asterisks indicate the levels of significant differences between samples imuno-precipitated by IgG and anti-GFP (Student’s t-test: *, P<0.05). **(C)** Schematic diagrams represented effector and reporter constructs used in the transient dual-LUC assays. The effector constructs contained the NaWRKY70 coding sequence driven by the CaMV35S promoter. The reporter construct contained the luciferase (LUC) driven by the promoter of NaF6’H1 and the Renilla luciferase (REN) driven by the CaMV 35S promoter. **(D)** In the transient dual-LUC assays, over-expression NaWRKY70 could significantly lead to the activation of NaF6’H1 promoter. Asterisks indicate the levels of significant differences between control and NaWRKY70 over-expression samples (Student’s t-test: *, P<0.05).

We next confirmed the binding of NaWRKY70 to *NaF6’H1* promoter by chromatin immune-precipitation-qPCR (Chip-qPCR) experiment in NaWRKY70-eGFP over-expressed transgenic lines. Higher levels of *NaF6’H* promoter regions around probe 1 were enriched with GFP antibody, indicating that NaWRKY70 could bind to the promoter of *NaF6’H1* via this W-box sequence *in vivo* (Figure 5B).

NaWRKY70 also activated *NaF6’H1* promoter transiently in dual-LUC system assay. As shown in Figure 5C and 5D, a significantly higher LUC/REN ratio was detected, which indicated that NaWRKY70 activated *NaF6’H1* promoter thereby enhanced LUC activity.

Thus, our data clearly indicates that NaWRKY70 binds to *NaF6’H1* promoter directly, and activates *NaF6’H1* expression.

### NaEBF2 and NaJAZs interact with NaWRKY70 to suppress *NaF6’H1* expression

To investigate the mechanism how NaWRKY70 integrates JA and ethylene signals, we constructed a yeast library with mixed leave samples of 1, 2 and 3 dpi, and screened proteins that interact with NaWRKY70. A total of 50 colonies were obtained in two-round of screening with high stringency selection medium. Among them, four were related to JA and ethylene signaling pathways, including NaEBF2, NaEIN3-like 1, NaJAZe1 and NaMYC2b.

EBF, an ethylene binding factor, is an important repressor in ethylene signaling pathway (Potuschak *et al*., 2003). There are three EBFs in *N. attenuata*. Yeast-two-hybrid experiments indicated that two NaEBFs could bind to NaWRKY70, they are NaEBF1 and NaEBF2 (Figure 6A, Supplementary Fig S8A). The interaction of NaEBF2 and NaWRKY70 was also confirmed by BiFC (Figure 6B). *NaEBF2* expression was elicited by *A. alternata* (Figure 6C). When *NaEBF2* was silenced via VIGS, *A. alternata*-induced scopoletin and scopolin levels were enhanced, and plants became more resistance to the fungus (Figure 6D-F). However, silencing *NaEBF1* did not affect scopoletin and scopolin production and plant resistance (Supplementary Fig. S8B-E). Interestingly, when *NaEBF2* and *NaWRKY70* were transiently co-overexpressed in leaves of *N*. *attenuata*, the endogenous expression of *NaF6’H1*, which was usually induced by NaWRKY70, was suppressed to control levels by NaEBF2 (Figure 6G). Furthermore, dual-LUC assay also showed that co-overexpression of *NaEBF2* and *NaWRKY70* suppressed the LUC/REN ratio driven by *NaF6’H1* promoter compare with those elicited by *NaWRKY70*-overexpression alone (Figure 6H and 6I). These results explain why treatment with MeJA alone cannot elicit *NaF6’H1* expression, because the induction of *NaF6’H1* by NaWRKY70 is still repressed by NaEBF2.

**Figure 6.**
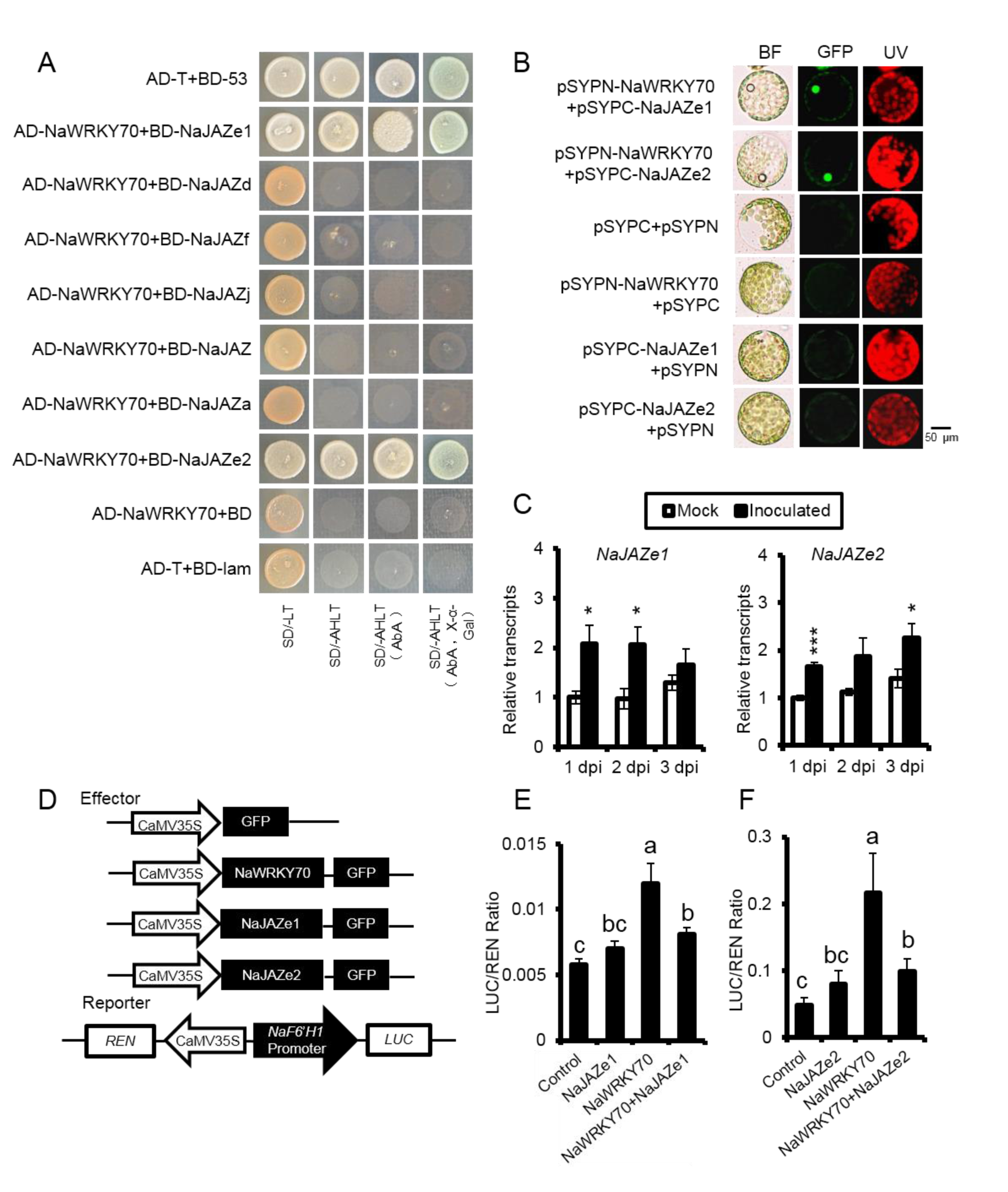
NaEBF2 interacts with NaWRKY70 to suppress the transcriptional activity of NaF6’H1. **(A)** Yeast-two-hybrid indicated the interaction of NaWRKY70 with NaEBF2. The yeast cells co-transformed with pGADT7-NaWRKY70 and pGBDT7-NaEBF2 were grown well on the tetra-dropout medium (SD/-Leu/-Trp/-His/-Ade, applying AbA and X-α-gal). Cells co-transformed with pGADT7-T and pGBDT7-53 were used as positive controls, and cells transformed with pGADT7-NaWRKY70 and pGBDT7, pGADT7 and pGBDT7-NaEBF2, pGADT7-T and pGBDT7-lam were used as negative controls. **(B)** Bimolecular fluorescence complementation (BiFC) showed the interaction between NaWRKY70 and NaEBF2 in *N. attenuata* protoplasts. The NaWRKY70 was fused at N-terminal of YFP (pSYPN-NaWRKY70), and NaEBF2 was inserted at C-terminal of YFP (pSYPC-NaEBF2). Protoplasts transformed with pSYPN+pSYPC, pSYPN-NaWRKY70+pSYPC or pSYPC-NaEBF2+pSYPN were used as negative controls. **(C)** Mean (±SE) relative *A. al*ternata-induced NaEBF2 expression levels were quantified by RT-qPCR at 1, 2 and 3 dpi. Asterisks indicate the level of significant differences between mock and inoculated samples (Student’s t-test: **, P<0.01; ***, P<0.005). Mean (± SE) relative NaEBF2 transcripts **(D)** and scopoletin and scopolin levels **(E)** were measured in 10 replicated young leaves of EV and VIGS NaEBF2 plants at 1 dpi. Asterisks indicate levels of significant differences between EV and VIGS plants (Student’s t-test: ***, P<0.005). **(F)** Mean (± SE) diameter of necrotic lesions of 20 biological replicates of young leaves of EV and VIGS NaEBF2 plants infected with A. alternata for 5 d (left). Photos of 6 representative leaves of each different genotype at 5 dpi were taken (right). Asterisks indicate levels of significant differences between EV and VIGS plants (Student’s t-test: ***, P<0.005). **(G)** Transient expression of both NaWRKY70 and NaEBF2 in N. attenuata leaves inhibited NaWRKY70-induced NaF6ʹH1 expression as quantified by RT-qPCR. Different letters indicate significant differences by one-way ANOVA followed by Duncan’s test (P<0.05). **(H)** Schematic diagrams represented effector and reporter constructs used in the transient dual-LUC assays. The effector constructs contained the NaWRKY70 or NaEBF2 coding sequence driven by the CaMV35S promoter. The reporter construct contained the luciferase (LUC) driven by the promoter of NaF6’H1 and the Renilla luciferase (REN) driven by the CaMV 35S promoter. **(I)** In the transient dual-LUC assays, NaWRKY70-overexpression led to the activation of NaF6’H1 promoter. However, NaEBF2 suppressed NaWRKY70-induced NaF6ʹH1 expression. Different letters indicate significant differences by two-way ANOVA followed by Duncan’s test (P<0.05).

Transcriptome analysis showed that seven NaJAZs were induced by *A. alternata* in the *N. attenuata* (Supplementary Table 2). However, only NaJAZe1 and NaJAZe2 were found to physically interact with NaWRKY70 protein by yeast two-hybrid (Figure 7A). These interactions were further confirmed by BiFC assay that the YFP fluorescence was observed when we co-expressed NaJAZe1-YFP^C^ and NaWRKY70-YFP^N^ or NaJAZe2-YFP^C^ and NaWRKY70-YFP^N^ constructs (Figure 7B). The expressions of *NaJAZe1* and *NaJAZe2* were also slightly induced by *A. alternata* (Figure 7C). As expected, the activation of *NaF6’H1* promoter by NaWRKY70 could be strongly suppressed when either *NaJAZe1* or *NaJAZe2* was co-expressed with *NaWRKY70* (Figure 7D-F). These results explain why treatment with ethephon alone cannot elicited *NaF6’H1* expression. Thus, our results reveal that three key repressors of JA and ethylene signaling pathways, NaJAZe1, NaJAZe2 and NaEBF2 physically interact with NaWRKY70, thereby strongly suppress NaWRKY70-regulated *NaF6’H1* expression. Our data uncovers the reason why treatment with MeJA or ethephon alone cannot elicited *NaF6’H1* expression.

**Figure 7.**
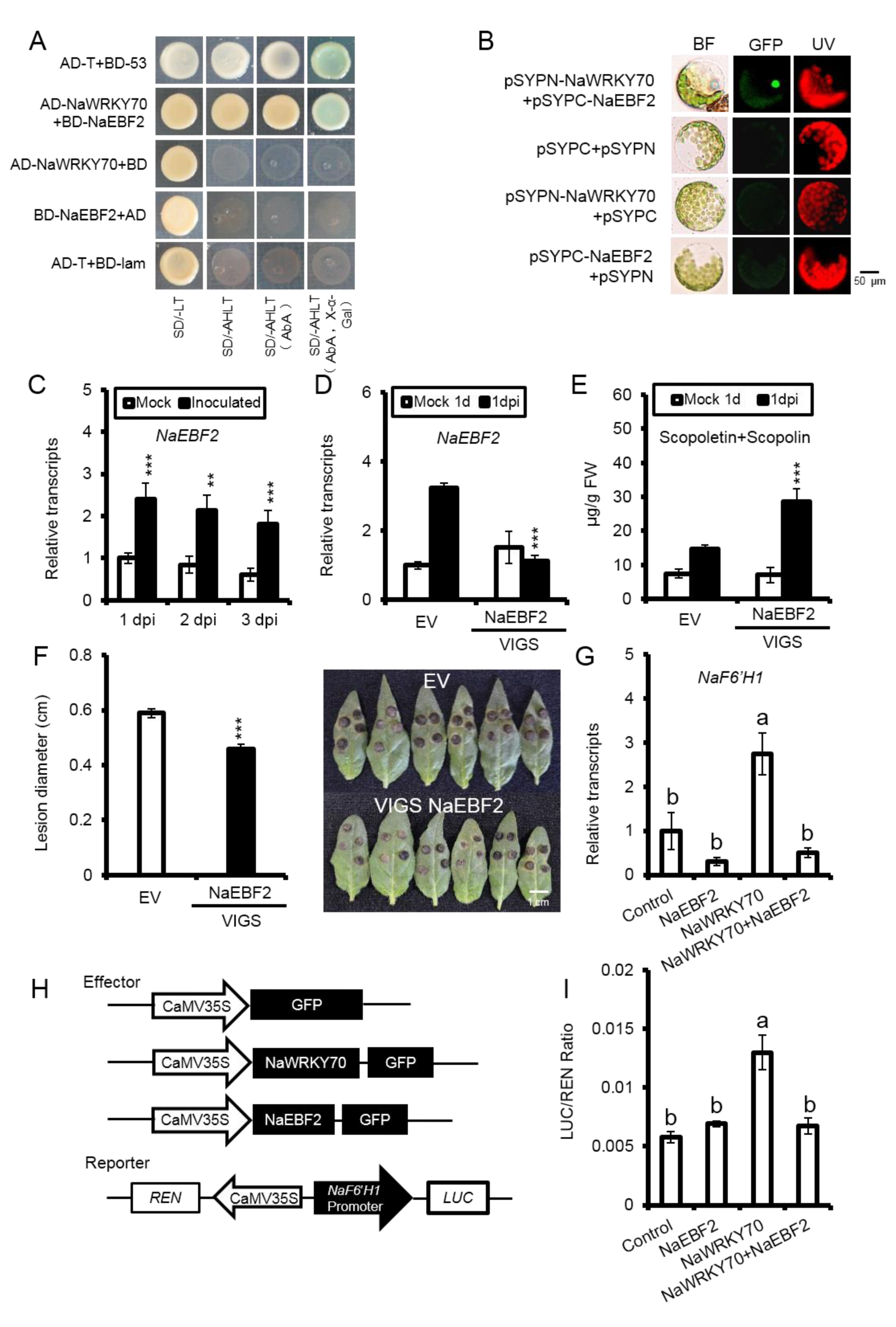
NaJAZe1 and NaJAZe2 interact with NaWRKY70 to inhibit the transcription activity of NaF6’H1. **(A)** Yeast two-hybrid indicated the interaction of NaWRKY70 with NaJAZs. The yeast cells co-transformed with pGADT7-NaWRKY70 and pGBDT7-NaJAZe1 or pGBDT7-NaJAZe1 were grown well on the dropout medium (SD/-Leu/-Trp/-His/-Ade, applying AbA and X-α-gal). However, yeast cells with pGADT7-NaWRKY70 co-transformed with pGBDT7-NaJAZd, pGBDT7-NaJAZf, pGBDT7-NaJAZj, pGBDT7-NaJAZ, or pGBDT7-NaJAZa could not grow on tetra-dropout mediums. pGADT7-T and pGBDT7-53 was used as positive controls, pGADT7-T and pGBDT7-lam were used as negative controls. **(B)** Bimolecular fluorescence complementation (BiFC) analysis showed the interaction between NaWRKY70 and NaJAZe1 or NaJAZe2 in protoplasts of N. attenuata. The NaWRKY70 was fused at N-terminal of YFP (pSYPN-NaWRKY70), and NaJAZe1or NaJAZe2 was inserted at C-terminal of YFP (pSYPC-NaEBF2). pSYPN+pSYPC, pSYPN-NaWRKY70+pSYPC, pSYPC-NaJAZe1+pSYPN and pSYPC-NaJAZe2+pSYPN were used as negative controls. **(C)** Mean (±SE) relative A. alternata-induced NaJAZe1 and NaJAZe2 expression levels were quantified by RT-qPCR at 1, 2 and 3 dpi. Asterisks indicate the level of significant differences between mock and inoculated samples (Student’s t-test: **, P<0.01; ***, P<0.005). **(D)** Schematic diagrams represented effector and reporter constructs used in the transient dual-LUC assays. The effector constructs contained the NaWRKY70, NaJAZe1 or NaJAZe2 coding sequence driven by the CaMV35S promoter. The reporter construct contained the luciferase (LUC) driven by the promoter of NaF6’H1 and the Renilla luciferase (REN) driven by the CaMV 35S promoter. In the transient dual-LUC assays **(E and F)**, NaWRKY70-overexpression led to the activation of NaF6’H1 promoter. However, co-transformed with NaWRKY70 and NaJAZe1 or NaJAZe2 could strongly suppress this transactivation. Different letters indicate significant differences by two-way ANOVA followed by Duncan’s test (P<0.05)

### NaEIN3-like1, a direct transcription regulator of *NaWRKY70*, interacts with NaWRKY70 to enhance *NaF6’H1* expression

To further confirm the protein interaction of NaWRKY70 and NaEIN3-like1 from yeast-two-hybrid screening, we further examined their interaction by using full-length cDNA sequences with bimolecular fluorescence complementation (BiFC) and yeast two-hybrid system. The yeast cells co-transformed with pGADT7-NaWRKY70 and pGBDT7-NaEIN3-like1 were grow well on the tetra-dropout medium (without Leu, Trp, His and Ade, applying AbA and X-α-gal) compare to negative controls (Figure 8A). In the BiFC assays, NaEIN3-like1 and NaWRKY70 were fused to the N-terminal and C-terminal fragments of YFP, respectively. Strong fluorescence signal was observed in protoplasts of *N. attenuata* when co-transforming with both NaEIN3-like1-YFP^N^ and NaWRKY70-YFP^C^, while no fluorescence signal were detected in all negative controls (Figure 8B). During dual-LUC reporter assay, *NaWRKY70* over-expression lead to the activation of *NaF6’H1* promoter, and this transactivation was even enhanced when *NaWRKY70* and *NaEIN3-like1* were both over-expressed (Figure 8C and 8D). We also transiently over-expressed *NaWRKY70* and *NaEIN3-like1* in *N. attenuata* leaves, and found that the levels of *NaF6’H1* expression activated by NaWRKY70 were enhanced when co-overexpressed with *NaEIN3-like1* (Figure 8E). EMSA results indicated that increasing amount of NaEIN3-like1 protein had not effect on the binding activities of NaWRKY70 to probes of *NaF6’H1* promoter (Figure 8F). These results suggested that NaEIN3-like1 interacts with NaWRKY70 to enhance *NaF6’H1* expression.

**Figure 8.**
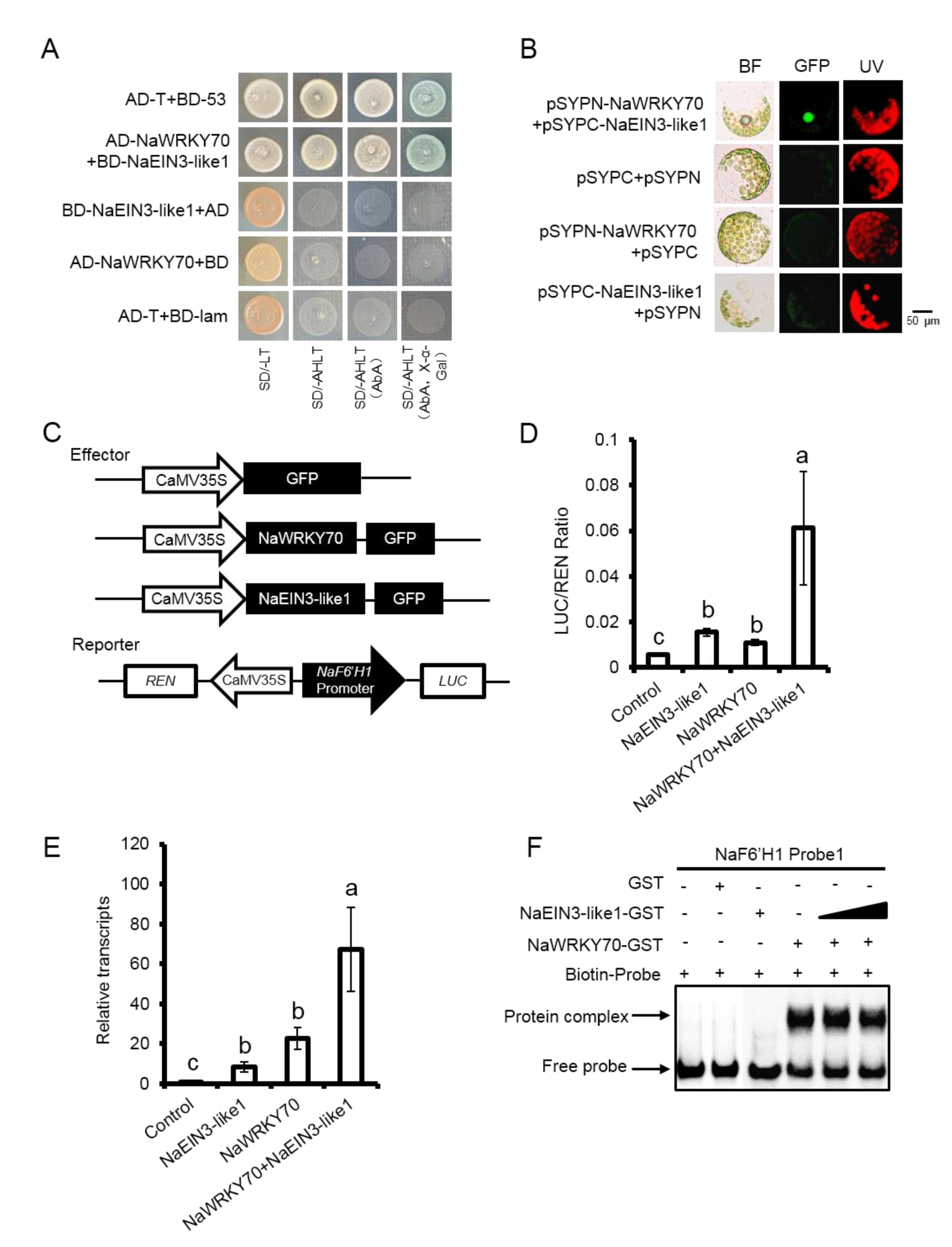
NaEIN3-like1 interacts with NaWRY70 to enhance NaF6’H1 expression. **(A)** Yeast two-hybrid indicated the interaction of NaWRKY70 with NaEIN3-like1. The yeast cells co-transformed with pGADT7-NaWRKY70 and pGBDT7-NaEIN3-like1 were grown well on the dropout medium (SD/-Leu/-Trp/-His/-Ade, applying AbA and X-α-gal). pGADT7-T and pGBDT7-53 was used as positive controls, pGADT7-NaWRKY70 and pGBDT7, pGADT7 and pGBDT7-NaEIN3-like1, pGADT7-T and pGBDT7-lam were used as negative controls. **(B)** Bimolecular fluorescence complementation (BiFC) analysis showed the interaction between NaWRKY70 and NaEIN3-like1 in protoplasts of N. attenuata. The NaWRKY70 was fused with N-terminal of YFP (pSYPN-NaWRKY70), and NaEIN3-like1 was inserted at C-terminal of YFP (pSYPC-NaEBF2). pSYPN+ pSYPC, pSYPN-NaWRKY70+pSYPC and pSYPC-NaEIN3-like1+pSYPN were used as negative controls. **(C)** Schematic diagrams represented effector and reporter constructs used in the transient dual-LUC assays. The effector constructs contained the NaWRKY70 or NaEIN3-like3 coding sequence driven by the CaMV35S promoter. The reporter construct contained the luciferase (LUC) driven by the promoter of NaF6’H1 and the Renilla luciferase (REN) driven by the CaMV 35S promoter. **(D)** In the transient dual-LUC assays, NaWRKY70-overexpression led to the activation of NaF6’H1 promoter. Co-expression of NaWRKY70 and NaEIN3-like1 could strongly enhance this transactivation of NaF6’H1 promoter. Different letters indicate significant differences by one-way ANOVA followed by Duncan’s test (P<0.05). **(E)** Co-transformation of NaWRKY70 and NaEIN3-like1 in N. attenuata leaves led to higher induction of NaF6ʹH1 expression as quantified by RT-qPCR. Different letters indicate significant differences by two-way ANOVA followed by Duncan’s test (P<0.05). **(F)** EMSA results indicated that increasing amount of NaEIN3-like1 protein had not effect on the binding activities of NaWRKY70 to probes of NaF6’H1 promoter.

When *NaEIN3-like1* was silenced by VIGS, *A. alternata*-elicited *NaF6’H1, NaWRKY70*, scopoletin and scopolin accumulation, and plant resistance to the fungus, were strongly impaired (Figure 9A-E). EMSA indicated that NaEIN3-like1 could specifically bind to one of the probes designed from *NaWRKY70* promoter, and this binding was abolished when 200 times of unlabeled or mutated probes were added (Figure 9F). To further analysis the regulatory effect of NaEIN3-like1 protein, we used a dual-LUC reporter system to explore the activation of NaEIN3-like1 protein on the transcriptional of *NaWRKY70*. After transforming 35s:: NaEIN3-like1 construct to *N. benthamiana*, the LUC/REN ratio were highly increased compare to the negative controls (Figure 9G and 9H), indicating NaEIN3-like1 could activate the LUC expression, which were under the control of *NaWRKY70* promoter.

**Figure 9.**
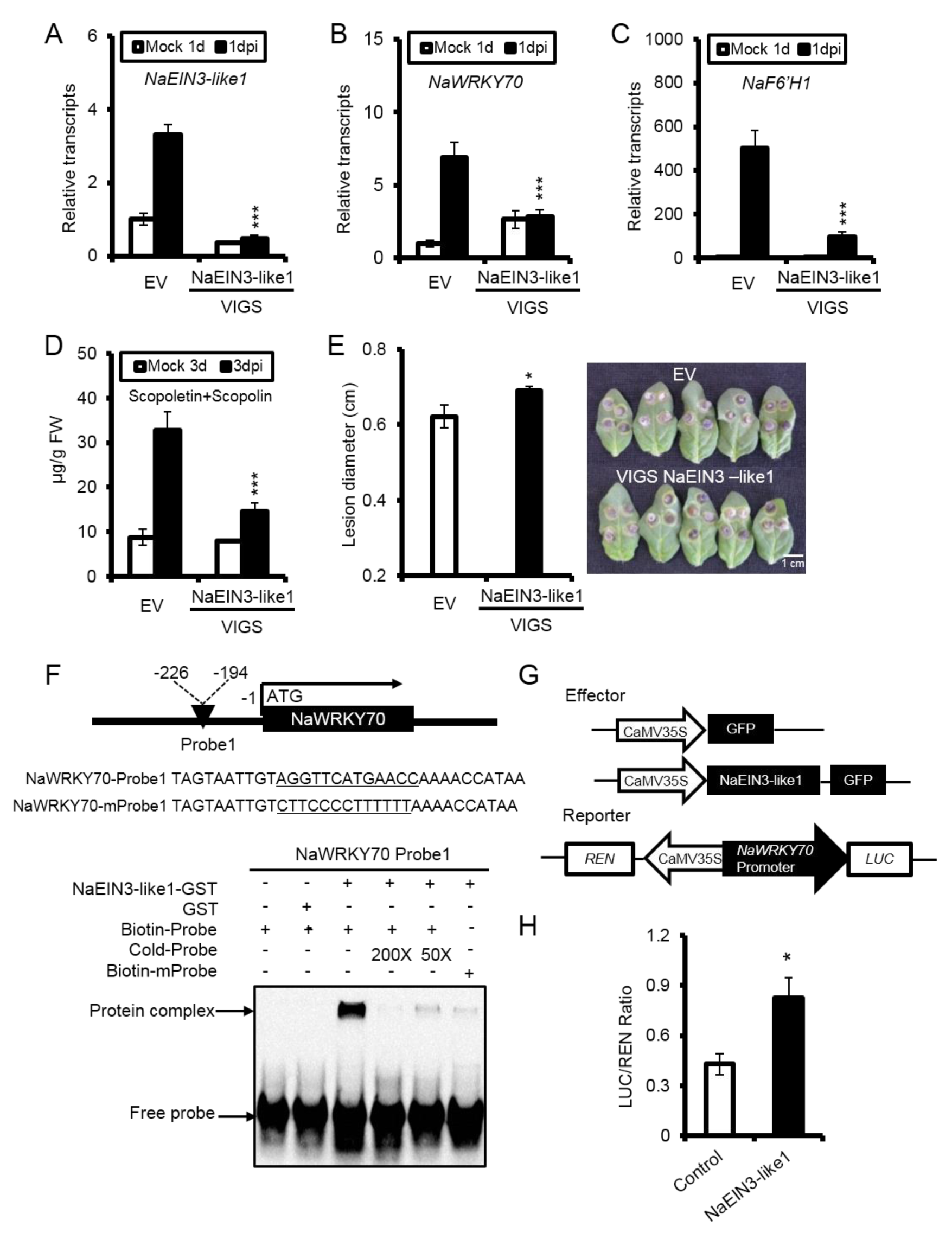
NaEIN3-like1 directly binds to the promoter of NaWRKY70 and activates its transcriptional activity. Mean (±SE) relative expression levels of *NaEIN3-like1* **(A)**, *NaWRKY70* **(B)** and *NaF6’H1* **(C)**, and scopoletin and scopolin levels **(D)** were measured in five biological replicated young leaves of EV and VIGS NaEIN3-like1 at 1 dpi. **(E)** Mean (± SE) diameter of necrotic lesions of 15 biological replicated rosette leaves of EV and VIGS NaEIN3-like1 infected with *A. alternata* for 5 d. Photos of 5 representative leaves of each different genotypes at 5 dpi were taken (right). Asterisks indicate the level of significant difference between WT and VIGS NaEIN3-like1 with the same treatments (Student’s t-test: *, P<0.05; ***, P<0.005). **(F)** Schematic diagrams of NaWRKY70 promoter and EMSA result of the binding of NaEIN3-like1 to NaWRKY70 promoter. The probe 1 (and mutated one: mProbe 1) was designed from NaWRKY70 promoter as indicated. EMSA demonstrated that NaEIN3-like1 protein could specifically bind to probe 1. The mobility shift was suppressed by cold unlabeled probes, and almost abolished when mutated probes were used. **(G)** Schematic diagrams represented effector and reporter constructs used in the transient dual-LUC assays. The effector construct contained the NaEIN3-like1 coding sequence driven by the CaMV35S promoter. The reporter construct contained the luciferase (LUC) driven by the promoter of NaWRKY70 and the Renilla luciferase (REN) driven by the CaMV 35S promoter. **(H)** In the transient dual-LUC assay, over-expression of NaEIN3-like1 could significantly lead to the activation of NaWRKY70 promoter. Asterisks indicate the levels of significant differences between control and NaEIN3-like1 over-expression samples (Student’s t-test: *, P<0.05).

Since NaEIN3-like 1 was identified to interact with NaWRKY70, and its expression was significantly elicited by *A. alternata*. We also checked other NaEIN3-like genes in transcriptome data of *A. alternata*-induced samples. We also found that another *NaEIN3*-like was induced at 1 dpi (Supplementary Table 2).This NaEIN3-like positively modulated *NaF6’H1* expression, scopoletin and scopolin biosynthesis, and plant resistance to *A. alternata* (Supplementary Fig. S9A-E).

However, this NaEIN3-like could not interact with NaWRKY70 in yeast, and could not bind to four predicted probes designed from *NaWRKY70* promote (Supplementary Fig. S9F and 9G), suggesting that this NaEIN3-like regulates *NaF6’H1* expression in some other unknown mechanisms.

Taken together, our findings uncover that 1) both NaEIN3-like and NaEIN3-like1 positively regulate *NaWRKY70* expression, and 2) at least NaEIN3-like1 directly binding and activating *NaWRKY70* expression, 3) NaEIN3-like1 interacts with NaWRKY70 to enhance *NaF6’H1*expression and scopoletin and scopolin biosynthesis.

### NaMYC2 interacts with WRKY70 to enhance *NaF6’H1* expression

Pervious study revealed the function of transcription factor MYC2 in JA signaling as a master regulator to modulate downstream target genes (Zhai and Li 2019; Chico *et al*., 2020). We silenced all NaMYC2 homolog genes (NaMYC2a, NaMYC2b, NaMYC2c and NaMYC2d, Supplementary Table 2) respectively in the *N. attenuata* via VIGS. Protein sequence analysis showed that NaMYC2a and NaMYC2b were highest similarity in clustered together (Supplementary Fig. S10). In VIGS NaMYC2a, VIGS NaMYC2b and VIGS NaMYC2a+2b plants, *A. alternata*-induced *NaF6’H1* expression, scopoletin, and resistance were strongly reduced (Figure 10A-D). In addition, *A. alternata*-induced transcripts of *NaF6’H1* was also impaired NaMYC2c-silenced plants, whereas they were not altered in VIGS NaMYC2d plants (Supplementary Fig. S11B and 11D). Interestingly, *A. alternata*-elicited *NaWRKY70* transcripts were decreased in VIGS NaMYC2a+2b and VIGS NaMYC2c plants (Supplementary Fig. S11A and 11C). Sequence analysis revealed that *NaF6’H1* promoter contained several G-boxes (CACGTG) or E-boxes (CANNTG). However, EMSA assays did not support the idea that NaMYC2a-GST, NaMYC2b-GST and NaMYC2d-GST proteins could bind to these boxes (Supplementary Fig. S11E-H).

**Figure 10.**
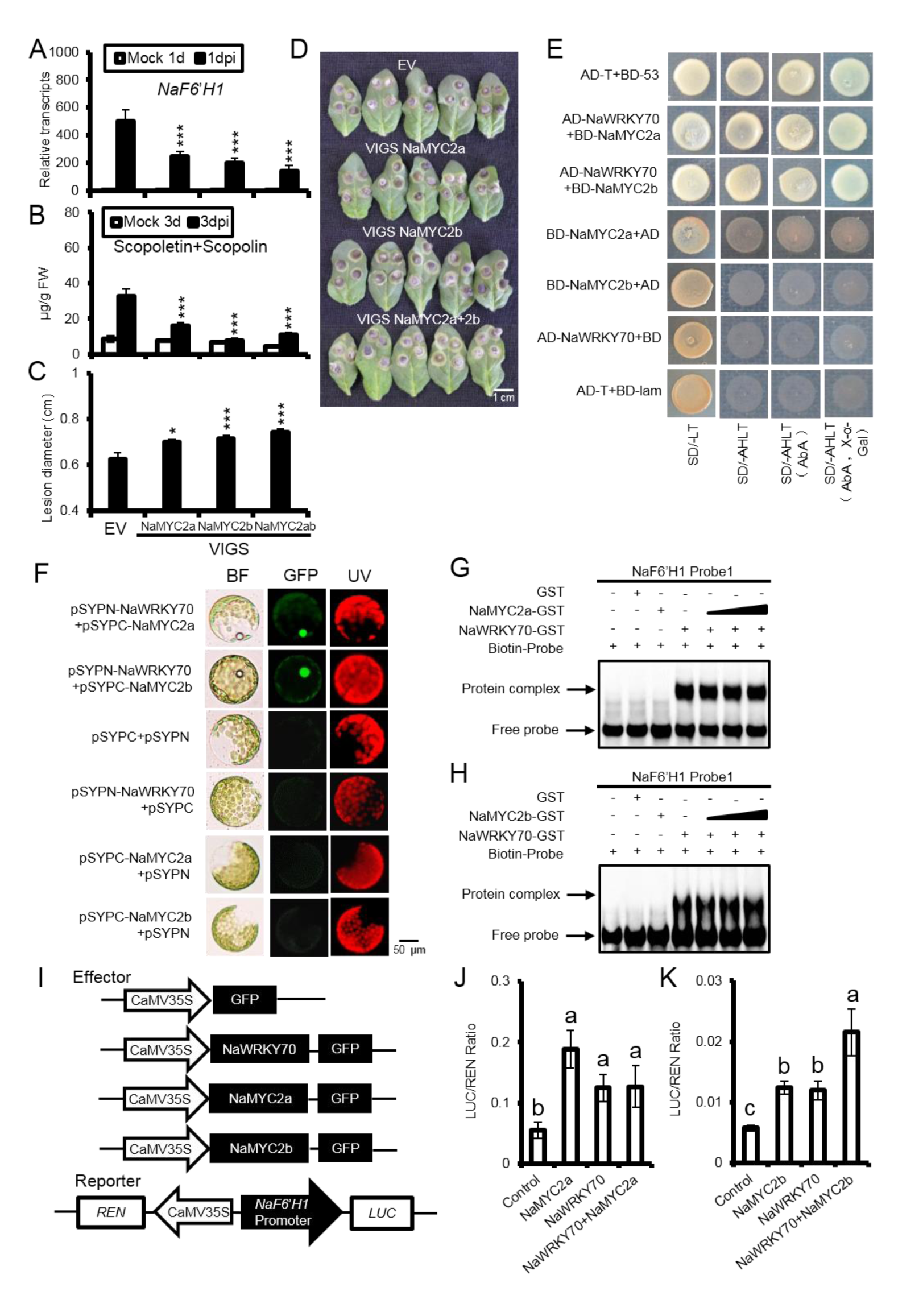
NaMYC2 interacts with WRKY70 to enhance NaF6’H1 expression. Mean (± SE) relative NaF6’H1 transcripts **(A)** and scopoletin and scopolin levels **(B)** were measured in ten-biological replicates of young leaves of EV, VIGS NaMYC2, VIGSMYC2b and VIGS MYC2a+2b plants at 1 dpi. Asterisks indicate levels of significant differences between EV and VIGS plants (Student’s t-test: ***, P<0.005). **(C)** Mean (± SE) diameter of necrotic lesions of 15 biological replicated young leaves of EV, VIGS NaMYC2, VIGSMYC2b and VIGS MYC2a+2b infected with A. alternata for 5 d. Asterisks indicate the level of significant difference between EV and VIGS plants (Student’s t-test: *, P<0.05; ***, P<0.005). **(D)** Photos of 5 representative leaves of each different genotype at 5 dpi were taken (right). **(E)** Yeast two-hybrid indicated the interaction of NaWRKY70 with NaMYC2a and NaMYC2b. The yeast cells co-transformed with pGADT7-NaWRKY70 and pGBDT7-NaMYC2a or NaMYC2b were grown well on the dropout medium (SD/-Leu/-Trp/-His/-Ade, applying AbA and X-α-gal). pGADT7-T and pGBDT7-53 was used as positive controls, pGADT7-NaWRKY70 and pGBDT7, pGADT7 and pGBDT7-NaMYC2a, pGADT7 and pGBDT7-NaMYC2b, pGADT7-T and pGBDT7-lam were used as negative controls. **(F)** Bimolecular fluorescence complementation (BiFC) analysis showed the interaction between NaWRKY70 and NaMYC2a or NaMYC2b in protoplasts of N. attenuata. The NaWRKY70 was fused with N-terminal of YFP (pSYPN-NaWRKY70), and NaMYC2a or NaMYc2b was inserted at the C-terminal of YFP (pSYPC-NaEBF2). pSYPN+pSYPC, pSYPN-NaWRKY70+pSYPC, pSYPC-NaMYC2a+pSYPN and pSYPC-NaMYC2b+pSYPN were used as negative controls. **(G and H)** EMSA results indicated that increasing amount of NaMYC2a-GST or NaMYC2b-GST proteins had not effect on the binding activities of NaWRKY70 to probes of NaF6’H1 promoter. **(I)** Schematic diagrams represented effector and reporter constructs used in the transient dual-LUC assays. The effector constructs contained the NaWRKY70, NaMYC2a or NaMYc2b coding sequence driven by the CaMV35S promoter. The reporter construct contained the luciferase (LUC) driven by the promoter of NaF6’H1 and the Renilla luciferase (REN) driven by the CaMV 35S promoter. **(J)** In dual-LUC assays, NaWRKY70 over-expression led to the activation of NaF6’H1 promoter. NaMYC2a could not enhance NaWRKY70-regulated NaF6’H1 promoter activities. **(K)** Both NaWRKY70 and NaMYC2b could induced NaF6’H1 promoter activities individually. Co-transformation of NaMYC2b and NaWRKY70 enhanced NaF6’H1 promoter activities. Different letters indicate significant differences by two-way ANOVA followed by Duncan’s test (P<0.05).

Yeast-two-hybrid confirmed the interaction of NaMYC2b and NaWRKY70 (Figure 10E). In addition, we also found that NaMYC2a could bind to NaWRKY70 (Figure 10E). These interactions were further confirmed by BiFC assay (Figure 10F). With the increasing amount of NaMYC2a-GST or NaMYC2b-GST proteins, the mobility shift signals of NaWRKY70 with *NaF6’H1* promoter probes were not altered, indicating the physical interaction of NaMYC2 to NaWRKY70 had not influence on the binding of NaWRKY70 to probes of *NaF6’H1* promoter (Figure 10G and 10H). The dual-LUC assay revealed that NaMYC2b but not NaMYC2a could enhance NaWRKY70-regulated *NaF6’H1* promoter activities (Figure 10I-K).

These results suggest that NaMYC2a, NaMYC2b and NaMYC2c are required for *A. alternata*-induced *NaWRKY70* and *NaF6’H1*exprssion, and importantly NaMYC2b could physically interact with NaWRKY70 to improve the transcriptional ability of *NaF6’H1*.

### Silencing *NaWRKY70* strongly reduces JA-Ile biosynthesis

Although *A. alternata*-induced *NaWRKY70* is dependent on both JA and ethylene signaling, it is currently unclear whether silencing *NaWRKY70* had any effect on JA and ethylene biosynthesis. We thus checked the transcriptional levels of *13-lipoxygenase* (*NaLOX3*), *allene oxide synthase* (*NaAOS*), *allene oxide cyclase* (*NaAOC*), *12-oxophytodienoate reductase* (*NaOPR*), *acyl-CoA oxidase* (*NaACX*), *peroxisomal 3-keto-acyl-CoA thiolase* (*NaKAT*), *jasmonic acid-amido synthetase* (*NaJAR*) and *cytochrome P450 94B3-like* (*NaCYP94B3*), all of which were shown to encode crucial enzymes of JA biosynthesis and metabolism respectively. Expression of *NaAOS*, *NaAOC*, *NaOPR2*, *NaOPR3*, *NaACX2*, *NaACX4*, *NaKAT5*, *NaJAR4* and *NaJAR6* was induced but at much lower levels in NaWRKY70-RNAi plants compared to those of WT in response to *A. alternata* at 3 dpi; while *NaLOX*3, *NaACX1*, *NaKAT1*, *NaJAR1* and *NaCYP94B3* were unchanged; and *NaACX2* expression showed increased (Figure 11B-D, Supplementary Fig. S12). Consistently, JA-Ile productions in two NaWRKY70-RNAi plants were strongly decreased, with only one-third of those of WT at 3 dpi (Figure 11A).

**Figure 11.**
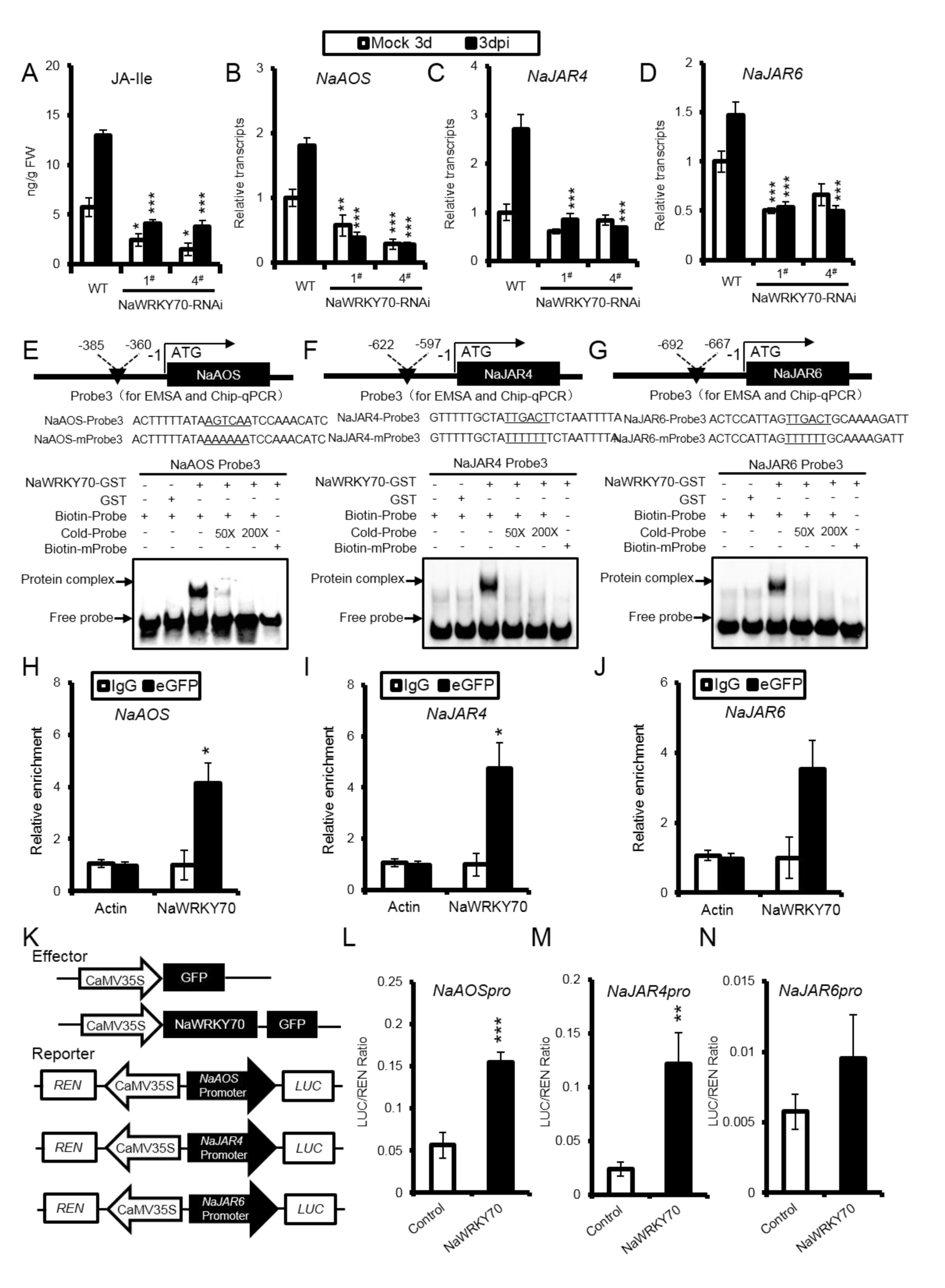
NaWRKY70 directly binds to the promoters of NaAOS, NaJAR4 and NaJAR6, and activates their expression. **(A)** Mean (±SE) A. alternata-induced JA-Ile levels were determined by HPLC-MS/MS in six biological replicates of 0 leaves of WT and two independent NaWRKY70-silenced plants (NaWRKY70-RNAi-1# and 4#) at 3 dpi. Mean (±SE) relative transcriptional levels of NaAOS **(B)**, NaJAR4 **(C)** and NaJAR6 **(D)** were measured by RT-qPCR in five biological replicates of WT and NaWRKY70-RNAi line 1# and 4# at 3 dpi. Asterisks indicate the levels of significant differences between WT and two NaWRKY70-RNAi lines with the same treatments (Student’s t-test: **, P<0.01; ***, P<0.005). EMSA assays demonstrated that NaWRKY70 protein could bind to promoters of NaAOS **(E)**, NaJAR4 **(F)** and NaJAR6 **(G)** in vitro. Biotin-labeled probes and mutant probes were designed from W-boxes as indicated in each gene. Chip-qPCR experiments indicated that NaWRKY70 could directly bind to the promoters of NaAOS **(H)**, NaJAR4 **(I)** and NaJAR6 **(J)** in vivo. Asterisks indicate the levels of significant differences between samples imuno-precipitated by IgG and anti-GFP (Student’s t-test: *, P<0.05). **(K)** Schematic diagrams represented effector (NaWRKY70 and control GFP) and reporter constructs used in the transient dual-LUC assays. The effector construct contained the NaWRKY70 coding sequence driven by CaMV35S promoter. The reporter constructs contained the luciferase (LUC) driven by promoters of NaAOS, NaJAR4 or NaJAR6 and the Renilla luciferase (REN) driven by the CaMV 35S promoter. In the transient dual-LUC assays, NaWRKY70 activates transcriptional activities of NaAOS **(L)**, NaJAR4 **(M)** and NaJAR6 promoters **(N)**.

As levels of *A. alternata*-induced transcripts of most biosynthetic enzyme genes and JA-Ile were strongly impaired in NaWRKY70-RNAi plants, we analyzed the W-box domains in the promoters of *NaAOS*, *NaOPR2*, *NaACX4*, *NaKAT5*, *NaJAR4* and *NaJAR6*. EMSA, dual-LUC assay and Chip-qPCR experiments all indicated that NaWRKY70 could directly bind to the promoters of *NaAOS*, *NaJAR4* and *NaJAR6 in vitro* and *in vivo*, and activate their transcriptional ability respectively (Figure 11E-N). Furthermore, we also found that NaWRKY70 could directly bind to the promoters of *NaOPR2*, *NaACX4* and *NaKAT5* by EMSA (Supplementary Fig. S13).

In addition, we investigated transcriptional levels of 1-aminocyclopropane-1-carboxylate synthase (NaACS1), 1-aminocyclopropane-1-carboxylate oxidase1 (NaACO1) and 1-aminocyclopropane-1-carboxylate oxidase2 (NaACO2), three crucial enzyme genes for ethylene biosynthesis. However, expression levels of all three genes and ethylene production were not altered in NaWRKY70-silenced plants (Supplementary Fig. S14).

Taken together, NaWRKY70 controls JA-Ile biosynthesis by binding to the promoters of biosynthetic enzymes as a feedback, but does not affect ethylene production when inoculated with *A. alternata*.

## Discussion

### Synergistic induction of scopoletin and scopolin biosynthesis by JA and ethylene signals

Phytoalexins are important defense arsenals produced in plants after pathogen attack, and are thus under the control of multi-interconnecting network of signaling pathways. Camalexin, a major phytoalexin in *Arabidopsis*, is regulated cooperatively through CPK5/CPK6 and MPK3/MPK6-mediated phosphorylation after *Botrytis cinerea* infection (Zhou *et al*., 2020). In maize, the accumulation of kauralexins and zealexins were induced synergistically by JA and ethylene co-treatments (Fu *et al*., 2018). We previously showed that *A. alternata*-elicited scopoletin and scopolin biosynthesis were dependent on JA or ethylene signaling pathways in *N. attenuata* (Sun *et al*., 2014b; Sun *et al*., 2017), while the regulatory mechanisms by these two signaling pathways remained unknown.

In this study, we demonstrated the synergistic regulation of scopoletin and scopolin biosynthesis by JA and ethylene signals. Scopoletin and scopolin were dramatically induced in *N. attenuata* leaves after co-treatments with MeJA and ethephon, but not by MeJA or ethephon alone (Figure 1). This synergistic induction was conserved in the genus of *Nicotiana* (Supplementary Fig. S2), and can be observed also in the lamina of young and mature leaves, mid-rib of young and mature leaves (Figure 1 and Supplementary Fig. S1). Importantly, the synergistic induction were abolished in irCOI1 (JA-insensitive) or Ov-etr1 (ethylene insensitive) plants (Supplementary Fig. S3), suggesting intact endogenous JA and ethylene signaling pathways were required. These results revealed that *A. alternata*-induced scopoletin and scopolin biosynthesis is regulated by JA and ethylene signals in a synergistic way.

### NaWRKY70 is required for synergistic induction of phytoalexins by JA and ethylene signals and *A. alternata* resistance

It has been reported that a series of transcription factors including WRKY, bHLH, MYB and ERF are played critical roles in the regulation of phytoalexins biosynthesis to defend pathogens attack. During the rice-Magnaporthe oryzae interaction, diterpenoid phytoalexins (DPs) biosynthesis were controlled by two transcription factors, WRKY45 and bHLH (DPF) (Akagi et al., 2014; Yamamura et al., 2015). GmMYB29A2, as a key regulator of phytoalexins glyceollin, directly influence its biosynthesis genes and metabolites in resistance to Phytophthora sojae (Jahan et al., 2020).

In this study, we demonstrated that NaWRKY70 plays an essential role in the synergistic induction of scopoletin and scopolin by JA and ethylene signals during *A. alternata* infection. By combination of RNA-sequencing, co-expression analysis and VIGS experiments, we identified a WRKY transcription factor NaWRKY70, whose expression was synergistically induced by MeJA and ethephon similar to *NaF6’H1* (Supplementary Fig. S4 and Figure 2). Plants silenced with *NaWRKY70* via RNAi or VIGS, and *NaWRKY70*-knockout plants generated by CRISPR/Cas9, were all highly susceptible to *A. alternata*, which were associated with strongly reduced levels of *NaF6’H1* and *NaCCoAOMTs* expression and scopoletin and scopolin production (Figure 3A-D; Figure 4A-C; Supplementary Fig. S6). Consistently, *NaWRKY70* over-expression plants exhibited highly *NaF6’H1* transcripts and scopoletin and scopolin levels and increased resistance to *A. alternata* (Figure 4D-G).

Notably, the synergistic induction of *NaF6’H1* expression, and production of scopoletin and scopolin, were strongly impaired in *NaWRKY70*-silenced plants after co-treatments of MeJA and ethephon (Figure 3E-G). Furthermore, EMSA, Chip-qPCR and dual-LUC assays further demonstrated that NaWRKY70 positively regulated *NaF6’H1* expression by direct binding and activating *NaF6’H1* promoter (Figure 5), confirming the important role of NaWRKY70 in regulation of *NaF6’H1* expression.

### NaWRKY70 is associated with age-dependent susceptibility to *A. alternata*

In many plant-pathogen interaction systems, the resistance against pathogens is usually dependent on the developmental age when host plants are infected. Some plants are more susceptible to diseases in early developmental stage and become more resistance when they are mature, for example rice and tobacco plants are more susceptible to *Xanthomonas oryzae* and *Phytophthora nicotianae* respectively in seedling stage. However, *Nicotiana* plants are more resistant to *A. alternata* in seedling stage while became susceptible when they are mature (Cheng and Sun, 2001; Sun *et al*., 2014a; Sun *et al*., 2014b). The mechanism behind this age-dependent susceptible to *A. alternata* is not currently clear. Previously, we have shown that young source-sink transition 0 leaves usually accumulated higher levels of JA, capsidiol, and scopoletin and scopolin than those of mature +3 leaves (Sun *et al*., 2014a; Sun *et al*., 2014b; Song *et al*., 2019). Here, we also found that the pattern of *A. alternata*-induced *NaWRKY70* were very similar to those of *NaF6’H1* expression and scopoletin and scopolin. They were all highly elicited in young leaves but this induction was decreasing while leaves became mature (Figure 2E-H). Considering the facts that NaWRKY70 functions as a key node of convergence for JA and ethylene signaling in scopoletin biosynthesis, and it also regulated *A. alternata*-induced JA-Ile level, we speculated that NaWRKY70 is a potential regulatory node of age and plant defense response to *A. alternata* in *Nicotiana* species.

### Feedback regulatory loop of NaWRKY70 in response to JA signaling

Interestingly, *A. alternata*-induced JA-Ile levels were dramatically decreased in *NaWRKY70*-silenced plants. Further real-time PCR results indicated that expressions of many JA biosynthetic and metabolism genes, including *NaAOS*, *NaAOC*, *NaOPR2*, *NaOPR3*, *NaACX2*, *NaACX4*, *NaKAT5*, *NaJAR4* and *NaJAR6* (Figure 11; Supplementary Fig. S12), were significantly reduced. EMSA and Chip-qPCR results revealed that NaWRKY70 could directly bind to the promoters of *NaAOS*, *NaJAR4* and *NaJAR6* and active their expression (Figure 11). Meanwhile, silencing *NaWRKY70* did not influence ethylene emissions and the transcripts of *NaACS1*, *NaACO1 and NaACO2* (Supplementary Fig. S14). Thus, NaWRKY70 plays a vital role in feed-backing de novo synthesis of JA-Ile by directly binding and activating multiple JA biosynthetic genes. Similarly, over-expression of *ZmWRKY79* enhanced levels of *ZmAOS1*, *ZmLOX1*, and *ZmJAZ17* (Fu *et al*., 2018). The feedback regulation of phytohormone synthesis by transcription factors seems to be a biological evolution consequence to quickly response to multiples stimulus, and amplify signaling (Ren *et al*., 2018; Zhang *et al*., 2019; Chu *et al*., 2021).

### NaWRKY70 integrates signals from JA and ethylene pathways in scopoletin and scopolin biosynthesis

Although JA and ethylene signals are usually associated with plant defense against necrotrophic pathogens and insect herbivores, complicated mode of interactions between JA and ethylene has been reported. For example, ethylene strongly suppresses JA-induced nicotine biosynthesis in *Nicotiana* species (Shoji *et al*., 2000). However, JA and ethylene also act synergistically in elicitation of phytoalexins kauralexins and zealexins in maize (Fu *et al*., 2018), and scopoletin and scopolin in *Nicotiana* species in this study. However, the mechanism behind their synergistic regulation by JA and ethylene is still unclear.

In *Arabidopsis*, *PDF1.2* is the best known defense gene synergistically regulated by JA and ethylene signals through ERF1 (Lorenzo *et al*., 2003). Later, JAZ proteins had been shown to interact with EIN3/EIL1 to suppress their regulation of *ERF1* expression. Thus, EIN3/EIL1 has been considered as the molecular link for the synergistic induction of *PDF1.2* by JA and ethylene signals. JA enhances the transcriptional regulation of *ERF1* by EIN3/EIL1 through removal of JAZs, while ethylene stimulates *EIN3*/*EIL1* expression and increase their protein levels (Zhu *et al*., 2011). In this study, we found a novel mechanism of synergistic induction of scopoletin and scopolin production by JA and ethylene signals through NaWRKY70.

Here, we found that NaEIN3-like1 not only regulates *NaWRKY70* expression directly, but also physically interacts with NaWRKY70 to enhance *NaF6’H1* expression. Thus, we demonstrate a new role of EIN3-like proteins in the synergistic induction of defense responses in *N. attenuata*. It acts both as an upstream transcriptional regulator and a protein interaction partner of NaWRKY70 to control scopoletin and scopolin biosynthesis.

EBF1 and EBF2 are EIN3 binding F-box proteins. They overlapped their function on stability of EIN3 as repressors in ethylene signaling (Yang *et al*., 2015; Zhao *et al*., 2021). In previous studies, EBFs are involved in various physiological processes including fruit ripening (Ding *et al*., 2019), seed germination (Gagne *et al*., 2004) and in response to high salinity in plants (Peng *et al*., 2014). However, the role of EBFs in synergistic induction of defense responses has not been reported. In this study, the expression of *NaEBF2* was significantly induced by *A. alternata*. Silencing *NaEBF2* enhanced scopoletin and scopolin biosynthesis and resistance against *A. alternata*. Importantly, yeast-two-hybrid and BiFC indicated the physical interaction of NaEBF2 and NaWRKY70. This interaction strongly suppressed the transcriptional activity of *NaF6’H1* (Figure 6). Thus, our results uncovered an important role of EBF in synergistic induction of scopoletin and scopolin by JA and ethylene signals, through physical interaction with NaWRKY70 to inhibit *NaF6’H1* expression.

In our previous study, we showed that NaMYC2a was involved in scopoletin and scopolin biosynthesis (Sun *et al*., 2014b). Here, we found that *A*. *alternata*-induced transcripts of *NaWRKY70* and *NaF6’H1*, and scopoletin and scopolin levels were significantly blocked in VIGS NaMYC2a, VIGS NaMYC2b and VIGS NaMYC2c plants except in VIGS NaMYC2d (Figure 10; Supplementary Fig. S11).We further showed that NaMYC2a, NaMYC2b and NaMYC2c protein could not likely directly bind to GCC- or E-boxes in *NaWRKY70* promoter (Supplementary Fig. S11). More importantly, we found that NaMYC2b interacted with NaWRKY70 in yeast-two-hybrid and BiFC assay. The interaction of NaMYC2b and NaWRKY70 enhanced the transcriptional activities of *NaF6’H1* (Figure 10). These results indicated that NaMYC2b augmented *NaF6’H1* expression and scopoletin and scopolin biosynthesis through interacting with NaWRKY70, and also positively regulated *NaWRKY70* expression.

Jasmonate ZIM Domain (JAZ) proteins are suppressors responsible for JA signaling pathway (Thines *et al*., 2007; Katsir *et al*., 2008). It has been reported that JAZ interact with TFs to participate in multiple physiological processes. For example, the interaction JAZ with complex protein WD-Repeat/bHLH/MYB to regulate anthocyanin accumulation in *Arabidopsis thaliana* (Qi *et al*., 2011); MdJAZ1 and MdJAZ2 are combining with MdBBX37 to repress the interactional function between MdBBX37 and MdICE1, which protein complex is negatively participate in cold tolerance in apple (An *et al*., 2021). Here, by screening NaJAZe1, NaJAZd, NaJAZf, NaJAZj, NaJAZ, NaJAZa and NaJAZe2, we identified that NaJAZe1 and NaJAZe2 could physically interact with NaWRKY70 (Figure 7). Importantly, the interactions between NaJAZe1 or NaJAZe2 and NaWRKY70 could suppress the transcriptional activity of *NaF6’H1*, thus negatively regulating scopoletin and scopolin biosynthesis (Figure 7).

Taken all together, our data uncovers a novel but complicate regulation network of scopoletin biosynthesis by JA and ethylene signaling pathways in a synergistic way. We demonstrate that NaWRKY70 is a key regulation node integrating signals from JA and ethylene signals, through transcriptional regulation of *NaWRKY70* and protein interaction with NaWRKY70, and finally controls *NaF6’H1* expression and phytoalexin biosynthesis (Figure 12). Without JA and ethylene signals, NaJAZe1, NaJAZe2 and NaEBF2, acting like locks, interact with NaWRKY70 to suppress *NaF6’H1* expression. However, during *A. alternata* infection or co-treatments with MeJA and ethephon, both ethylene and JA signals are activated. The inhibition of NaWRKY70 by NaJAZe1, NaJAZe2 and NaEBF2 are released. NaEIN3-like and NaEIN3-like 1 will up-regulate *NaWRKY70* expression, and especially NaEIN3-like1 will not only directly bind to *NaWRKY70* promoter, activate *NaWRKY70* expression, but also physically bind to NaWRKY70 to enhance *NaF6’H1* expression. Similarly, NaMYC2b also regulates *NaWRKY70* expression, and interacts with NaWRKY70 protein to enhance *NaF6’H1* expression. Thus, our findings provide a perfect example of defense responses synergistically regulated by JA and ethylene signaling pathways, and extend our understandings of defense responses regulated by phytohormones in *Nicotiana* species to *A. alternata*.

**Figure 12.**
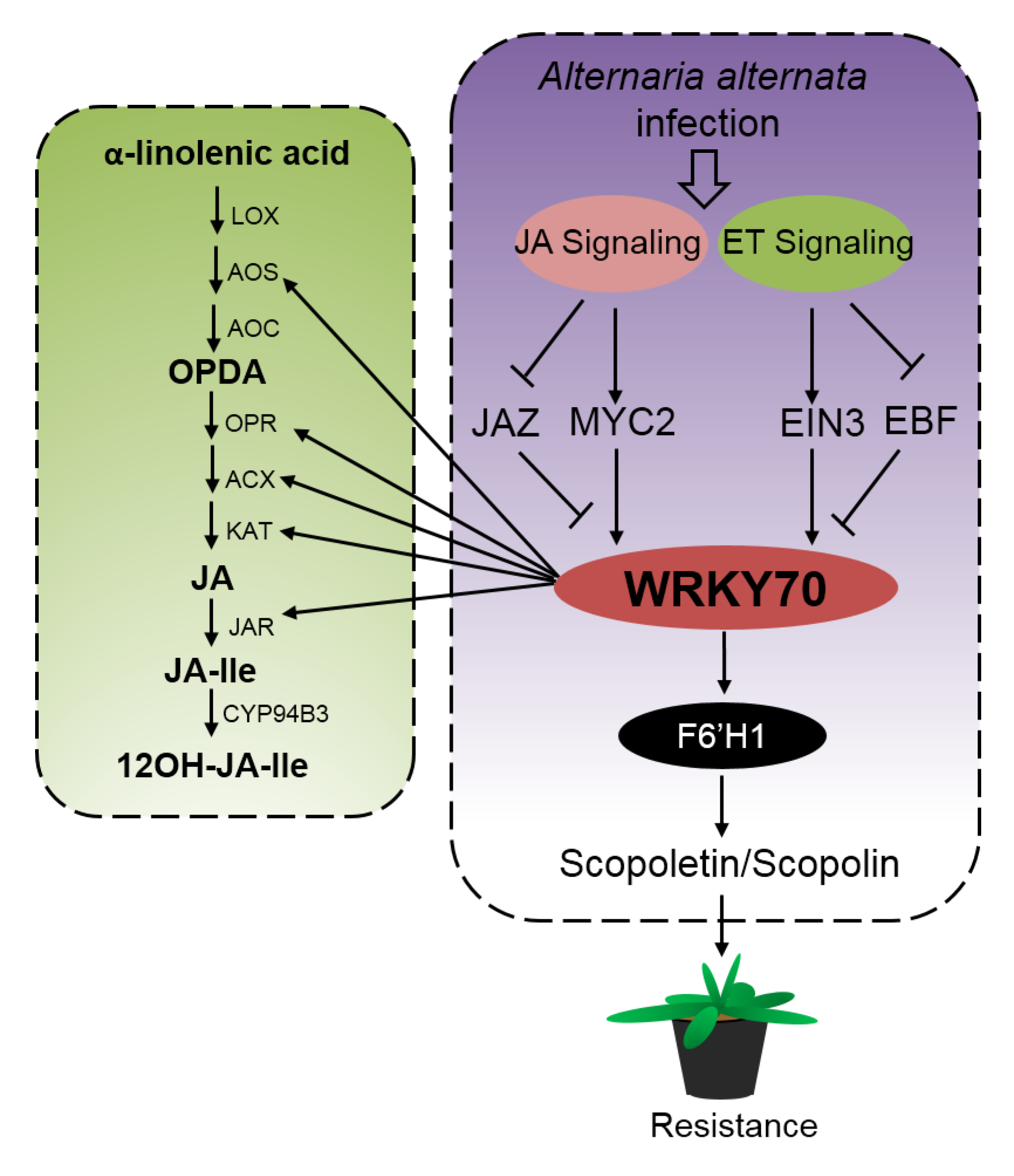
The working model of integration of both JA and ethylene signals by NaWRKY70 in defense responses against A. alternata in N. attenuata. In N. attenuata plants, both JA and ethylene signaling pathways are activated by A. alternata, and subsequently control the biosynthesis of phytoalexins scopoletin and its β-glycoside form, scopolin, and expression of their key enzyme gene NaF6’H synergistically. NaWRKY70 is an important node of this synergistic regulation; it binds and activates downstream target gene NaF6’H1 directly, and integrates upstream JA and ethylene signaling to enhance or inhibit NaF6’H1 expression. Several JA and ethylene signaling elements play a vital role in modulation of NaF6’H1 expression through NaWRKY70. NaJAZe1, NaJAZe2 and NaEBF2 proteins, acting as negative regulation factors of JA and ethylene signaling, interact with NaWRKY70 to suppress NaF6’H1 expression, which consequently inhibits scopoletin and scopolin accumulation without activation of both JA and ethylene signaling. On the other hand, NaMYC2b and NaEIN3-like1 are positive regulation factors of JA and ethylene; both regulate NaWRKY70 expression, and interact with NaWRYK70 to enhance NaF6’H1 expression. In addition, NaWRKY70 directly regulates A. alternata-induced JA-Ile levels by activating NaAOS, NaOPR2, NaACX4, NaKAT5, NaJAR4 and NaJAR6 genes as a feedback.

## Materials and Methods

### Plant and fungal materials

Seeds of 35^th^ generation of an *N. attenuata* inbred line was used as wild-type (WT) genotype. The seeds of *N. attenuata* transgenic lines including irACO (ethylene reduced; irACO), Ov-etr1 (ethylene insensitive), irAOC (deficient in JA biosynthesis) and irCOI1 (JA-insensitive) plants were generated previously (von Dahl *et al*., 2007; Kallenbach *et al*., 2012; Paschold *et al*., 2007), and provided by Prof. Ian T. Baldwin (Max-Planck institute of Chemical Ecology). Seed germination and plant growth were conducted as described by Krügel *et al*. (2002). *Alternaria alternata* were grown and inoculated on leaves as described by Sun *et al*. (2014a).

### Hormone Treatment

The source-sink transition leaves of *N. attenuata* plants (32-day-old) were sprayed with 1 mM methyl jasmonate (MeJA, Sigma) or 5 mM ethephon (2-chloroethanephosphonic acid, an ethylene-releasing agent, Sigma). Both MeJA and ethephon were prepared with distilled water. The leaves treated with H2O were served as a negative control. Samples were harvested at the appropriate times for further analysis.

### RNA extract and quantitative real-time PCR (RT-qPCR) assay

Total RNA was isolated by TRI reagent (Invitrogen) and cDNA was synthesized as described by Wu *et al*. (2013). Quantitative real-time PCR (qRT-PCR) was performed using the CFX Connect qPCR device (Bio-Rad) with iTaq™ Universal SYBR Green Supermix (Bio-Rad) and specific primers of genes (Supplementary Table S1) according to the manufacturer’s instructions. A linear standard curve (obtained from threshold cycle number versus log DNA quantity) was obtained by using a series of dilution of a specific cDNA sample, and the transcriptional levels of unknown samples were calculated according to this standard curve (Song *et al*., 2019). *NaActin2* gene whose expression was not altered by MeJA, ethephon and *A. alternata* treatments (Xu *et al*., 2018) was used as an internal standard.

### RNA-seq data analysis

After treated 6 h with water control, MeJA (1 mM), ethephon (5 mM) and co-treatment of MeJA and ethephon, three biological replicates of WT source-sink transition leaves with the same treatments were mixed for RNA preparation. RNA sequencing was conducted by Shanghai OE-Biotech (http://www.oebiotech.com/) with the Illumina Hiseq 2000. Sequencing was performed at 8 G depth, and mapped to the *N. attenuata* reference genome sequence. The differential expression between water control and MeJA and ethephon treatment samples with log2 (fold change)≥3 and its significance were calculated.

The raw sequence data reported in this paper has been deposited in the Genome Sequence Archive in BIG Data Center, Beijing Institute of Genomics (BIG), Chinese Academy of Sciences, under accession numbers CRAxxxxxx (waiting for the deposit number).

### Generation of virus-induced gene silencing plants

The highly specific fragment of the genes, including NaMYC4-like, NaMYC2-like, NaMYB24-like, NaMYB57-like, NaHAT5-like, NabHLH18-like, NaERF17-like, NaZinc655-like, NaWRKY70, NaMYB4-like, NaWRKY75-like, NaWRKY70-like, NaWRKY41-like, NabHLH137-like, NabHLH93-like, NaMYB44-like, NaEIN3-like, NaEIN3-like1, NaMYC2a, NaMYC2b, NaMYC2c, NaEBF1 and NaEBF2 were amplified by specific primer pairs respectively (Supplementary Table 1). The PCR fragments were digested with Hind III and Bam HI, and cloned into pTV00 vector. The recombinant vectors were introduced into *Agrobacterium tumefaciens* GV3101 strain (Saedler and Baldwin, 2004). The empty vector pTV00 (EV) and pTV00 vector carrying the phytoene desaturase (PDS) were used as controls respectively (Saedler and Baldwin 2004). When *PDS*-silenced plants occur with photo-bleaching phenotype, the young leaves of VIGS plants were harvested for measuring silencing efficiency. Around 20 biological replicates were used for each experiment.

### Generation of the NaWRKY70-RNAi, -knockout and over-expression plants

The highly specific inverted sequence of NaWRKY70 was amplified by primers (Supplementary Table S1) and inserted into the pRESC8 vector for RNAi. The full coding sequence of NaWRKY70 was amplified by primers (Supplementary Table S1) and inserted into the pCAMBIA1301-eGFP vector for over-expression. The recombinant vectors were introduced into *N. attenuata* plants by *Agrobacterium tumefaciens* LBA4404 strain via transformation as described by Zhao *et al*. (2020).

Single-insertion RNAi (NaWRKY70-RNAi-1^#^ and NaWRKY70-RNAi-4^#^) and over-expression lines (Ov-NaWRKY70-1^#^ and Ov-NaWRKY70-2^#^) were identified, bred to homozygosity in T2 generation and used in this study.

For further analysis the role of NaWRKY70, we obtained the NaWRKY70 knockout lines by CRISPR/Cas9 system. The guide RNA cloning vector pUC57-DT1T2 and Cas9/gRNA vector pHSE401 were constructed as described by Xing *et al* (2014). The 20 bp sequences o GCGGCCGAGAACTCACGCGC and GTCGGAGATGTATTGACTG were selected as the target site1 and sites2. The CRISPR/Cas9-NaWRKY70 vector was transformed into *Agrobacterium* strain LBA4404, and *Agrobacterium*-mediated transformation was performed as described by Liang and Wu (2022).

### Subcellular localization of NaWRKY70

The coding sequence of NaWRKY70 was cloned into the pM999 vector. The 35S::NaWRKY70-eGFP was transformed into the *N. attenuata* protoplasts. The *N. attenuata* protoplasts were prepared as described by Yoo *et al*. (2007) and Song *et al*. (2019). After incubation for 16-18 h, the signal of 35S:: NaWRKY70-eGPF were observed by using a fluorescent microscope (Leica DM5500 B). The empty vector pM999 was used as a negative control.

### Transient activation of target promoters by NaWRKY70

The promoters of *NaWRKY70*, *NaF6’H1, NaAOS, NaJAR4 and NaJAR6* were amplified by specific primers (Supplementary Table S1) and cloned into the pCAMBIA3301-LUC-REN vector reporters. The coding sequence of NaEIN3-like1, NaWRKY70, NaMYC2a, NaMYC2b, NaJAZe1, NaJAZe2 and NaEBF2 were amplified by specific primers (Supplementary Table S1) and insert into pCAMBIA3301 vector effectors. The constructed reporter and effector vectors were transformed into *Agrobacterium tumefaciens* GV1031 strain, and transiently expressed in *N. benthamiana* leaves. The luciferase (LUC) and renilla luciferase (REN) activity were measured according to dual Luciferase assay Kit (YEASEN).

The LUC/REN ratio were determined the transcriptional activity of the promoter. Supplementary Table S1 listed the primers used for these different genes. All experiments were performed at least five biological replicates.

To further analysis the effect of NaEIN3-like1, NaMYC2a, NaMYC2b, NaJAZe1, NaJAZe2 and NaEBF2 on NaWRKY70-*NaF6’H1* module, we also co-transformed these protein genes with *NaWRKY70* into *N. attenuata* leaves to measure *NaF6’H1* expression levels by RT-qPCR.

### Electrophoretic mobility shift (EMSA) assays

The full-length coding sequence of *NaWRKY70*, *NaEIN3-like*, *NaEIN3-like1*, *NaMYC2a*, *NaMYC2b* and *NaMYC2c* were amplified by specific primers (Supplementary Table S1), cloned into the pGEX-4T-1 vector fusing the glutathione S-transferase (GST) and transferred into the *E.coli* strain BL21. The fusion-proteins were induced by Isopropyl-beta-D-thiogalactopyranoside (IPTG, 0.01-0.05mM), then harvested the bacteria cells and purified with GST-tag purification resin (Beyotime). The detection of the binding of the recombinant protein and the probes labeled with biotin, were carried out with a chemiluminescent EMSA kit (Beyotime) according to the protocol suggested by the manufacture.

### Quantification of scopoletin and scopolin, JA-Ile and Ethylene

The leaf samples around 0.2 g were harvested and ground to fine powder in liquid nitrogen. The levels of scopoletin and scopolin were determined by HPLC-MS/MS as described in Sun *et al*. (2014b). JA-Ile were extracted and quantified by HPLC-MS/MS as described by Wu *et al*. (2008). Ethylene were extracted and quantified by HPLC-MS/MS as described by Zhang *et al*. (2021). **Yeast-two-hybrid assays**

The full-length coding sequence of NaWRKY70 was fused to pGADT7 to construct the bait vectors with EcoR I and Xho I. The full-length coding sequence of the *NaEIN3-like*, *NaEIN3-like1*, *NaMYC2a*, *NaMYC2b*, *NaEBF1*, *NaEBF2*, *NaJAZe1*, *NaJAZd*, *NaJAZf*, *NaJAZj*, *NaJAZ*, *NaJAZa* and *NaJAZe2* were introduced into the prey vector pGBDT7 via Nco I and Bam HI. These bait and prey vectors were co-transfected into yeast strain Y2H GOLD (Clontech). The transfected yeast cells were screened on SD/-Leu/-Trp, SD/-Ade/-His/-Leu/-Trp, SD/-Ade/-His/-Leu/-Trp (applying Aba) and SD/-Ade/-His/-Leu/-Trp (applying Aba and X-gal) medium cultured at 28°C for 5 d. Y2HGold (pGADT7-T/pGBDT7-lam) were used as negative controls and Y2HGold (pGADT7-T/pGBDT7-53) were used as positive controls. The primers used to amplify sequences were listed in Supplemental Table 1.

### Chromatin immune-precipitation assay

Chromatin immunoprecipitation was performed with EpiQuik Plant ChIP Kit (EPIGENTEK) according to the user manual. The source-sink transition leaves of over-expression NaWRKY70-eGFP plants (1.5-2 g) infected with *A. alternata* were prepared for ChIP assays. Relative enrichments were measured by RT-qPCR. All primers used in the ChIP experiments were listed in Supplemental Table 1.

Chromatins were precipitated with IgG and with GFP antibody but using primers detecting Actin served as the negative controls.

### Bimolecular fluorescent complementation (BiFC) assays

The full- length coding sequence of *NaWRKY70* was cloned into pSYP fusion with YFP N-terminal to generate pSYPN-NaWRKY70. The *NaEIN3-like1*, *NaMYC2a*, *NaMYC2b*, *NaEBF2*, *NaJAZe1* and *NaJAZe2* full-length CDS were inserted at the YFP C-terminal of pSYP vector, then co-transformed into *N. attenuata* protoplasts as described by Song *et al*. (2019). At 18-24 h after co-transformation, the YFP signals were observed under a fluorescent microscope (Leica DM5500 B). The primers used for BiFC were listed in Supplemental Table S1.

### Accession Numbers

The GenBank accession numbers of the genes in this article are as follows:

NaMYC4-like (XM_019377229.1), NaMYC2-like (NaMYC2d) (XM_019380820.1), NaMYB24-like (XM_019373720.1), NaMYB57-like (XM_019370259.1), NaHAT5-like (XM_019367916.1), NabHLH18-like (XM_019373104.1), NaERF17-like (XM_019378163.1), NaZinc655-like (XM_019390032.1), NaWRKY70 (XM_019399160.1), NaMYB4-like (XM_019391424.1), NaWRKY75-like (XM_019404314.1), NaWRKY70-like (XM_019372971.1), NaWRKY41-like (XM_019374936.1), NabHLH137-like (XM_019380008.1), NabHLH93-like (XM_019401728.1), NaMYB44-like (XM_019376996.1), NaEIN3-like (XM_019382253.1), NaEIN3-like1 (XM_019403482.1), NaMYC2a (XM_019398607.1), NaMYC2b (XM_019368208.1), NaMYC2c (XM_019375733.1), NaEBF1 (XM_019404974.1), NaEBF2 (XM_019374641.1), NaJAZe1(XM_019388454.1), NaJAZd (XM_019391388.1), NaJAZf (XM_019369600.1), NaJAZj (XM_019379894.1), NaJAZ (XM_019392390.1), NaJAZa (XM_019385384.1), NaJAZe2 (XM_019398932.1), NaLOX3 (XM_019399844.1), NaAOS (XM_019399851.1), NaAOC (XM_019406620.1), NaOPR2 (XM_019387447.1), NaOPR3 (XM_019400466.1), NaACX1 (XM_019372546.1), NaACX2 (XM_019408430.1), NaACX4 (XM_019383488.1), NaKAT1 (XM_019380608.), NaKAT3 (XM_019411655.1), NaKAT5 (XM_019385127.1), NaJAR1 (XM_019374932.1), NaJAR4 (XM_019383891.1), NaJAR6 (XM_019370360.1), NaCCoAOMT5 (XM_019376905.1), NaCCoAOMT6 (XM_019402596.1), NaACS1 (XM_019368410.1), NaACO1 (XM_019391407.1) and NaACO2 (XM_019393834.1).

### Author Contributions

Jinsong Wu and Na Song conceived the project, designed the experiments and prepared the manuscript; Na Song performed all experiments and analyzed the data.

### Supplement date

**Fig. S1.** Synergistic induction of scopoletin and scopolin in mid-rib of young leaves, lamina and mid-rib of mature leaves by MeJA and ethephon

**Fig. S2.** Synergistic induction of scopoletin and scopolin are absent in *Arabidopsis thaliana*, *Gossypium hirsutum,* and Solanaceae plants *Solanum tuberosum* and *Solanum lycopersicum*

**Fig. S3.** Requirements of intact endogenous JA and ethylene signaling pathways for the synergistic induction of scopoletin and scopolin by MeJA and ethephon

**Fig. S4.** *A. alternata*-elicited *NaF6’H1* expression as measured in plants transformed with EV or silenced with 16 candidate TFs individually via VIGS

**Fig. S5.** Alignment of protein sequences of NaWRKY70 and its homologs, and nuclear localization of NaWRKY70

**Fig. S6.** Silencing *NaWRKY70* impairs *A. alternata*-induced expressions of *NaCCoAOMTs*

**Fig. S7.** EMSA results indicates the binding of NaWRKY70 to probes designed from *NaF6’H1* promoter

**Fig. S8.** NaEBF1 interacts with NaWRKY70, but silencing *NaEBF1* does not affect *A. alternata*-induced scopoletin and scopolin production, and plant resistance

**Fig. S9.** NaEIN3-like does not interact with NaWRKY70 in yeast cells, but is required for *A. alternata*-induced transcripts of *NaWRKY70* and *NaF6’H1* and scopoletin and scopolin production

**Fig. S10.** Sequence alignment of NaMYC2a, NaMYC2b, NaMYC2c and NaMYC2d **Fig. S11.** NaMYC*2*s regulate *A. alternata*-induced *NaWRKY70* and *NaF6’H1* expressions indirectly

**Fig. S12.** Effect of silencing *NaWRKY70* on expressions of JA biosynthesis genes **Fig. S13.** NaWRKY70 directly binds to promoters of *NaOPR2*, *NaACX4* or *NaKAT5* **Fig. S14.** Silencing *NaWRKY70* has little impact on *A. alternata*-induced ethylene production and expression of ethylene biosynthetic genes

## Acknowledgments

We thank Prof. Ian T. Baldwin (Max-Planck Institute for Chemical Ecology, Jena, Germany) for providing irAOC, irCOI1, irACO and Ov-etr1 transgenic *N. attenuata* seeds and Biological Technology Open Platform of Kunming Institute of Botany for greenhouse and instrument services. This project was supported by the National Natural Science Foundation of China (NSFC Grant No. 31670262) and top-talent recruitment program of Yunnan to Prof. Jinsong Wu.

